# Molecular Configuration, Regulation and Function of Heterochannel Electrical Synapses

**DOI:** 10.1101/2025.09.14.675793

**Authors:** Atal Vats, Muraleedharan Sudhanand, Shivangi Verma, Ananya Bandyopadhyay, Nayantara Varma, Sandhya P Koushika, Vatsala Thirumalai, Abhishek Bhattacharya

## Abstract

Neuron-specific expression of particular gap junction channel components defines the configuration and functional properties of electrical synapses. However, how a neuron utilises multiple, simultaneously expressed channel proteins - connexins or innexins -to make meaningful connections with distinct synaptic partners remains largely unknown. Using the posterior mechanosensory circuit in *C. elegans,* we discovered that individual electrical synapses can be formed by clustering together molecularly distinct gap junction channel-types made of three different innexin proteins, INX-1, UNC-7, and UNC-9. In this previously unknown configuration, which we term as heterochannel synapses, molecularly distinct gap junction channel types functionally collaborate to regulate posterior touch sensory behaviour, enhancing functional robustness. We show that the synaptic trafficking of the molecularly different channel types within a heterochannel synapse is independently regulated by discrete and conserved kinesin motor proteins, while distinct molecular pathways involving channel-specific retrograde kinesins regulate their turnover. These independent, channel-specific regulations also make individual synapse-level alterations in the composition of heterochannel synapses possible under altered environmental conditions, providing a novel mechanism for electrical synapse plasticity. Finally, we present evidence of heterochannel electrical synapses in *C. elegans* locomotory circuits and in the cerebellar Purkinje neurons of zebrafish larvae. Altogether, we demonstrate a novel heterochannel organization of electrical synapses, their regulation, and functional importance, which may be a conserved feature of metazoan nervous system.

## INTRODUCTION

How a nervous system processes information and generates behaviour depends on synaptic connectivity and the molecular properties of synapses. While the role of molecular composition of chemical synapses has been widely explored, how the molecular compositions of the electrical synapses regulate circuit function and behaviour remains much understudied. Nonetheless, the significance of electrical synapses in nervous system development and function has been well established. For instance, fast signal transmission through electrical synapses has been shown to play multiple indispensable roles in nervous systems across species, including regulation of escape responses^1–7^.

The mechanosensory circuit responsible for low-threshold touch avoidance in *C. elegans* offers a tractable, well-mapped circuit to understand how the molecular composition of electrical synapses affects behaviour. Briefly, this circuit is wired by three anterior touch receptor neurons (TRNs) (ALML/R and AVM) and three posterior TRNs (PLML/R and PVM)^8^. Both the anterior and posterior set of TRNs form multiple chemical and electrical synapses with downstream interneurons, including premotor interneurons, governing locomotion^9–11^. Despite the well-characterized connectivity identified through electron micrograph (EM) studies, the specific functional contributions of these electrical synapses in touch sensation remain speculative.

Electrical synapses are formed bp analogous families: connexins in vertebrates and innexins in invertebrates (Figure S1A). It has been shown that individual neurons within vertebrate and invertebrate nervous systems often express multiple innexin or connexin genes in complex, combinatorial patterns ^12–14^. Although this availability of multiple connexins or innexins could theoretically allow for gap junctions with diverse molecular compositions (Figure S1A), recent evidences indicate that the availability of multiple connexin or innexin proteins do not necessarily result in the formation of multi-component gap junctions or synapse establishment^13,15,16^. The cellular mechanisms that determine the selection and utilization of available connexins or innexins for the establishment of synapses with precise specificity and functional properties remain to be fully understood. Similarly, little is known about the precise cellular mechanisms that govern synaptic trafficking of gap junction proteins, their assembly into specific electrical synapses, and their turnover. It also remains to be determined whether these cellular processes could be plastic. Understanding these regulatory principles will ultimately have significant bearings on the wiring and functional output of the connectome. This paper attempts to address these challenges.

We describe here that, unlike previously thought, even individual electrical synapses in the *C. elegans* nervous system can be formed by clustering molecularly-distinct gap junction channel-types, resulting in a “heterochannel” configuration. Focusing on posterior TRN, PLM, we show that individual electrical synapses are formed by clustering together three distinct gap junction channel types: INX-1 homotypic channels (made of INX-1 alone) and by reciprocally arranged UNC-7 and UNC-9 heterotypic channels. These molecularly distinct channel types within individual heterochannel synapses functionally collaborate to regulate PLM-mediated posterior low-threshold touch response. We further demonstrate that discrete channel-type-specific synaptic trafficking, as well as turnover mechanisms independently regulate dynamics of INX-1 and UNC-7/UNC-9 gap junction channels within individual heterochannel synapses in PLM neurons, thereby actively maintaining the molecular identity or the heterochannel-code of these synapses. Furthermore, in a remarkable example of synapse-specific plasticity, we demonstrate that the heterochannel-code of specific synapses in PLM gets reversibly altered under adverse environmental conditions when *C. elegans* larvae develop into the hibernation-like dauer diapause state^17–19^. Finally, we also provide evidences of functionally important heterochannel electrical synapses within locomotory circuits in *C. elegans* as well as their presence in the zebrafish brain, suggesting that this may be a fundamental design principle for electrical synapses across species. Taken together, our findings highlight the functional significance, regulation and plasticity of electrical synapses that are encoded at the level of the heterochannel code.

## RESULTS

### Individual electrical synapses in PLM neurons are composed of three distinct innexins

Low-threshold mechanical stimuli received at the posterior part of the *C. elegans* body are sensed by a pair of bilaterally symmetric Touch-Receptor Neurons (TRN), called PLML/R. Cell-ablation studies have highlighted that PLM neurons are absolutely important in this process^20^. We have also found that *unc-86(n846)* animals where TRNs fail to differentiate, were insensitive to posterior touch (Figure 1A). However, the specific roles of different synaptic connections of PLM in regulation of the mechanosensation still remain speculative. We investigated the roles of electrical synapses made by PLM neurons in touch elicited responses. Previous studies have shown that PLM neurons form electrical synapses to multiple downstream interneurons in two distinct synaptic zones^9,10,21^. At zone 1, proximal to the cell body, PLM forms electrical synapses with forward pre-motor interneuron, PVC and other interneurons, LUA and PVR (Figure S1B)^9,10^. At zone 2, at the distal end of the neuron, it synapses onto the BDU interneuron (Figure S1A)^21^. Previous cell ablation studies of the pre-motor interneurons have suggested a central role for the PLM-PVC electrical synapses in posterior touch sensation^20^. These electrical synapses have previously been shown to be formed by innexin, UNC-9^21,22^. To understand the potential role of electrical synapses in the posterior mechanosensation, we looked at the posterior touch sensory response in *unc-9(e101)* mutants. Surprisingly, we did not observe any posterior touch response defects associated with *unc-9(e101)* mutants (Figure 1A). Previous studies also corroborate that *unc-9* mutants do not significantly affect the posterior low-threshold mechanosensation in *C. elegans*^23,24^. In many instances, neurons in both vertebrate and invertebrate nervous systems have been shown to express multiple connexins or innexins, including PLM neurons^12,13,25^. Thus, we hypothesised that other PLM-innexins could play functionally redundant roles at electrical synapses in this circuit. To address this question, we performed a behavioural screen with all the innexins expressed in PLM and their electrical synaptic partners to identify mutants that led to a compromised posterior touch response. Our data showed that other than *inx-1*, which had a partial defect, none of the other innexins were required individually for posterior touch responses (Figure 1A). These results indeed suggested that there could be functional redundancy among PLM-expressed innexins. However, the inter-relationships between these innexins have not been characterised. Moreover, how an individual neuron utilizes its multiple connexin or innexin repertoire to build distinct electrical synapses with specific synaptic partners that may have distinct functional properties, has not been clearly understood.

**Figure 1:**
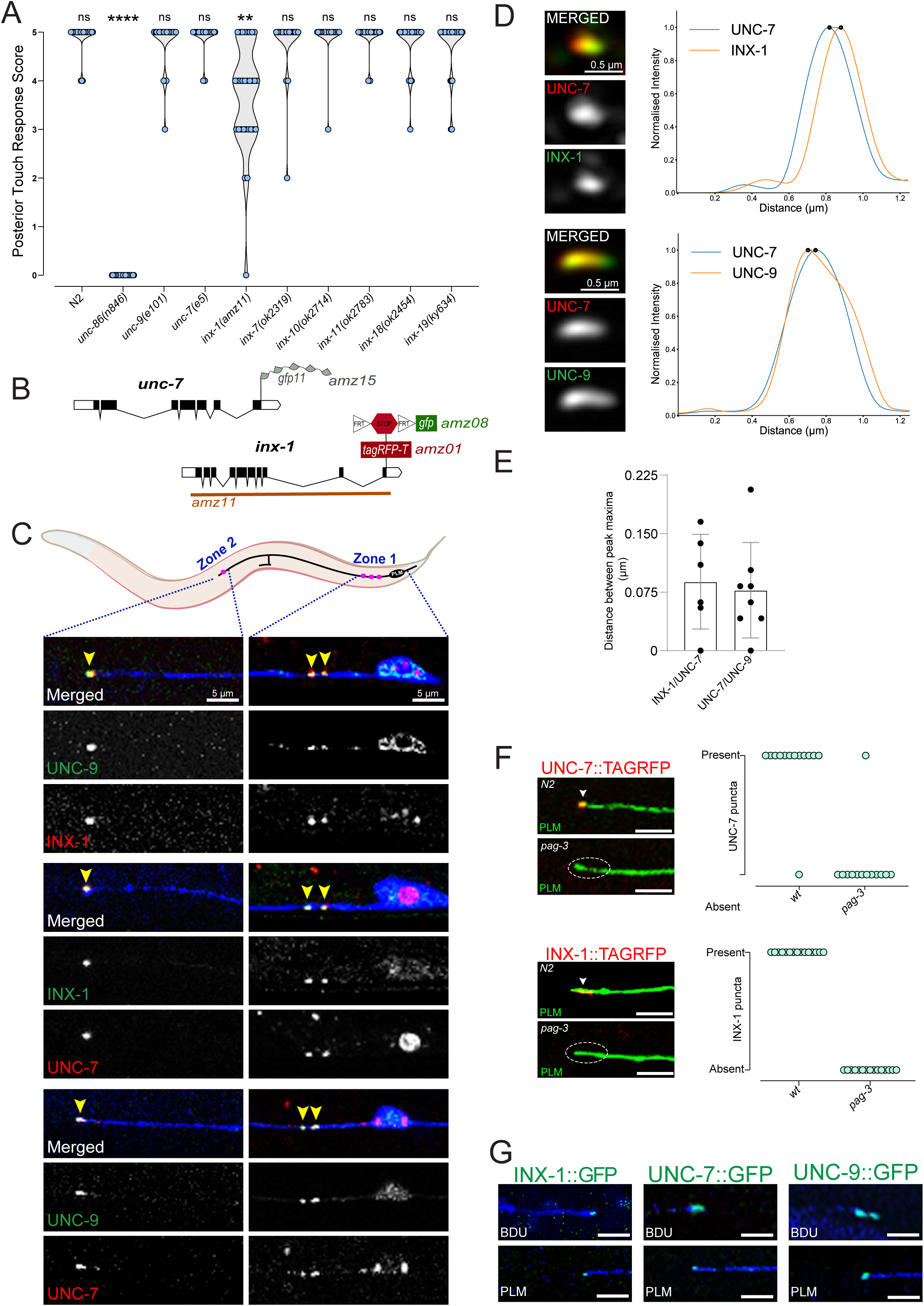
INX-1, UNC-7 and UNC-9 innexins co-localize in PLM electrical synapses. (A) Posterior low-threshold touch response in young adult animals. Each data point represents response of a single animal (n=30 animals/genotype). Statistical comparison between different groups were performed using the Fisher-Freeman-Halton exact test. n.s., non-significant, *p < 0.05, **p < 0.01, ***p < 0.001, ****p < 0.0001. Among innexin genes expressed in PLM neurons or in PLM-synaptic partners, only *inx-1(amz11)* mutants showed significant reduction in posterior touch response. *unc-86(n846)* animals were used as positive control. (B) Schematics showing 5Xsplit-GFP based *unc-*7 endogenous reporter allele*, unc-7(amz15[unc-7::5xgfp11])*, FLP/FRT-based cell-specific *inx-1::FLPon::gfp* reporter allele*, inx-1(amz08[inx-1::FRT::stop::FRT::egfp])*, TAGRFP-tagged reporter allele of *inx-1, inx-1(amz01[inx-1::tagrfp])* and deletion allele of *inx-1*, *inx-1(amz11)*. (C) Schematic of PLM neuron, showing the two distinct electrical synapse zones. Pairwise co-localization of endogenously-tagged (1) inx-1-*inx-1(amz01[inx-1::tagrfp]),* INX-1::EGFP(*amz08, amzEx79), (2) unc-7-* UNC-7::TAGRFP(*ot895)*, and UNC-9::7Xsplit-GFP(*miz81; amzEx112)* within PLM electrical synaptic puncta (yellow arrowheads) in both Zone-1 and Zone-2. PLM axons were marked by BFP expression (*amzIs01[mec-18p::ebfp2])*. Yellow asterisk marks *ectopic expression of mam-5p::tagrfp::h2b* co-injection marker in amzEx79. Scalebar = 5 μm. (D) SIM images showing co-localization of INX-1::EGFP(*amz08, amzEx79)* and UNC-7::TAGRFP(*ot895*), and UNC-7::TAGRFP(*ot895*) and GFP::UNC-9 (*amzEx76*), within PLM electrical synaptic punctum in zone-1. Corresponding plots show overlap of min-max normalized fluorescence intensity profiles across synaptic punctum. Black dots mark peak maxima for each curve. Scalebar = 500 nm. (E) Quantification of lateral distance between intensity-maxima peaks within a single punctum obtained in SIM for each innexin pair. Error bars indicate standard deviation. (F) Endogenously labelled UNC-7::TAGRFP(*ot895*) puncta was lost from the PLM zone-2 synapses in *pag-3(Is20)* mutant animals. Quantifications are shown on the right. (G) Endogenously labelled INX-1::TAGRFP(*amz01)* puncta was lost from the PLM zone-2 synapses in *pag-3(Is20)* mutant animals. Quantifications are shown on the right. (H) BDU- and PLM-specific expression of INX-1 (INX-1::FLPon::GFP) using cell-specific flippase-recombination showed punctate localization of INX-1 from both BDU and PLM partner-neuron sides (axons marked in blue) at the PLM synaptic zone-2. (I, J) BDU-and PLM-specific expression of endogenous UNC-7::GFP and UNC-9::GFP *using unc-7::5Xsplit-GFP* and *unc-9::7Xsplit-GFP* alleles, respectively, showed punctate localization of UNC-7 and UNC-9 from both BDU and PLM partner-neuron sides (axons marked in blue) at the PLM synaptic zone-2.

To understand how different innexins could participate in PLM-synapse formation and regulate the touch elicited responses, we set out to identify innexins that localize to the PLM electrical synapses in characteristic punctate patterns. We expressed GFP-or TagRFP-T-tagged individual innexins specifically in TRNs under a TRN-specific *mec-4* driver. We found that while UNC-9, INX-1 and UNC-7 showed electrical synapse-like stereotypic punctate localizations in both the electrical synapse zones in PLM (Figure S1B), other innexins (*inx-2, inx-7, inx-10, inx-18*) did not show punctate localizations at any of the synaptic zones (Figure S1A). We validated these observations by using endogenous GFP-or TagRFP-T-tagged reporter alleles for innexins representing both innexin cohorts. Using *inx-18(amz05[inx-18::TagRFP-T])* as a representative, we found that despite forming synaptic puncta in the head of the animal, endogenous INX-18 did not localize to the PLM synapses (Figure S1C). These results suggested that expression of an innexin is necessary, but may not be sufficient to drive synaptic localization.

To understand how INX-1, UNC-7 and UNC-9 might function in PLM electrical synapses, we first checked localization of endogenously GFP-or TagRFP-tagged reporter alleles of these innexins in pairwise manner. We used endogenously TagRFP-tagged reporter alleles of *inx-1*, *inx-1(amz01[inx-1::TagRFP-T])* and *unc-7*, *unc-7(ot895[unc-7::tagRFP-T])*^26^, and a split-GFP based reporter allele of *unc-9*, *unc-9(miz81[unc-9::7×gfp11])*^27^ (Figure 1B). Our results suggested that INX-1, UNC-7 and UNC-9 precisely colocalise in all PLM synapses (Figure 1C). Furthermore, we used super resolution, Structured Illumination Microscopy (SIM), to confirm that mean distances between the maximum intensity peaks of INX-1::GFP, UNC-7::TagRFP and UNC-9::GFP present within an individual PLM electrical synapse are very close (within 150nm), which indeed suggested colocalization of these three innexins at the PLM electrical synapses (Figure 1D).

Apart from their primary role in gap junctions, innexins are also known to function as undocked hemichannels^28–30^. Previously, it was shown that UNC-9 forms gap junction channels within PLM^21,22^. We went on to test whether INX-1 and UNC-7 also form bonafide gap junction channels in PLM. We reasoned that if INX-1 and UNC-7 are involved in formation of trans-synaptic gap junction channels, then loss of PLM-synaptic partner neuron should abrogate their punctate localization. To test this hypothesis, we observed the localization of INX-1 and UNC-7 puncta at the distal synapse of PLM formed with a single synaptic partner, BDU (Zone-2). We found that the punctate localizations of INX-1 and UNC-7 at the PLM-BDU synapses were completely abrogated in animals mutant for Zn-finger transcription factor, *pag-3(ls20), C. elegans* ortholog of Sensless/Gli, where BDU neurons fail to differentiate properly^31^ (Figure 1F, G). These results suggested that INX-1, UNC-7 and UNC-9 form gap junction channels at the PLM-electrical synapses.

Innexins can participate in gap junction formation from either or both synaptic partner sides. Moreover, all these innexins are expressed in both PLM and its synaptic partner neurons^12,13^, making it difficult to unambiguously answer this question. To this end, we cell-specifically labelled these innexins from PLM and corresponding partner neurons, and observed their localisation. We used flippase (FLP)-mediated cell-specific labelling strategy (Figure 1B) to endogenously label INX-1 with GFP, *inx-1(amz08[inx-1::FLPon::gfp])* in specific neuron-types. To cell-specifically label UNC-7, we generated a split-GFP-based^32^ reporter allele of *unc-7*, *unc-7(amz15[unc-7::5xgfp11]),* where a cassette containing five tandem repeats of 11^th^ beta-strand of GFP (*sfGFP11x5*) was inserted at the C-terminus of the *unc-7* locus (Figure 1B). Our data revealed that all three innexins-INX-1, UNC-7 and UNC-9 localise from both PLM and BDU synaptic sides at the Zone-2 synapse (Figure 1H, I, J). We also validated this dual-sided expression of INX-1 in the PLM-PVC synapse (Figure S1E). Overall, these results suggested that multiple innexins can be localised to individual electrical synapses, potentially attributing to the synapse function and ultimately to the posterior mechanosensory behaviour.

### Molecularly distinct channels cluster to form an electrical synapse

Innexins co-expressed in synaptic partners can interact with each other to form gap junction channels having distinct molecular configurations, ultimately dictating the functional properties of the channels^15,33–36^. To understand how INX-1, UNC-7 and UNC-9 could potentially regulate PLM-synaptic properties, we set out to test the composition of gap junctions made from these proteins. We hypothesised that INX-1, UNC-7 and UNC-9 proteins could either be forming independent, homotypic channels of their own or could be forming intermixed, heterotypic, or heteromeric channels with each other based on their oligomerisation and docking compatibilities. We reasoned that if two innexins interact to form a channel, the absence of one innexin should affect the punctate localisation of the partner innexin, i.e., innexins forming heteromeric or heterotypic channels should display mutual dependency, as shown for other innexins and connexins^13,33,37^. We observed that punctate localisation of endogenously tagged INX-1 remained unaffected in animals mutant for either *unc-7* or *unc-9,* as well as in unc-7, *unc-9* double mutant animals (Figure 2A). As gap junctions are trans-synaptic channels, it is still possible for INX-1 to interact in trans with other innexins expressed in the synaptic partners of PLM. To test this possibility, we curated a list of innexins that are expressed in PLM partner neurons^12,13^ (Figure S2A), and found that INX-1 localisation also remained unaffected in animals mutant for these innexins. We conclude that INX-1 is forming homotypic channels (Figure S2B). This is also consistent with our previous findings that INX-1 localizes from both synaptic partner sides in all PLM synapses (Figure 1H, S1D).

**Figure 2:**
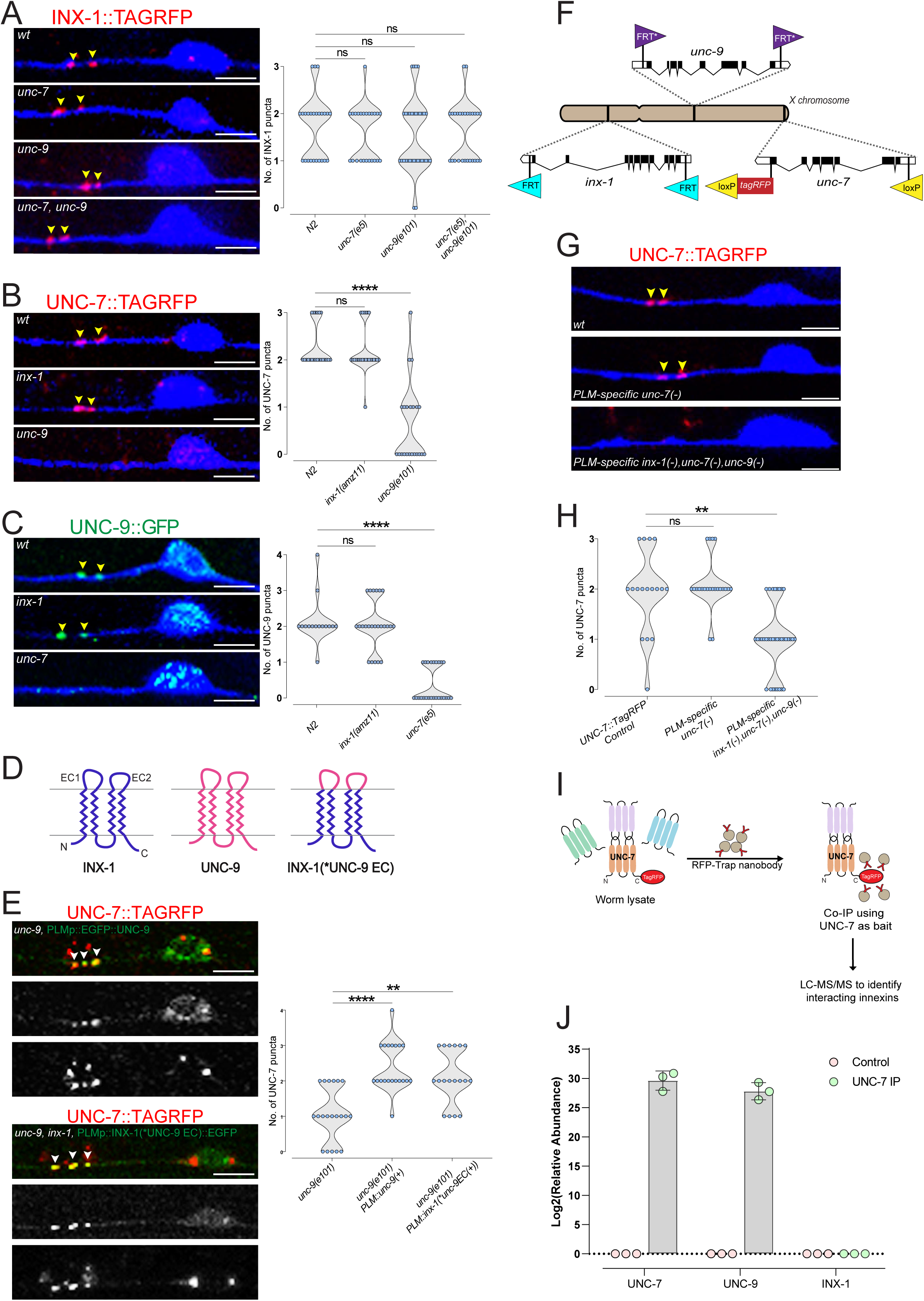
INX-1, UNC-7 and UNC-9 form molecularly heterogeneous gap junction channels within heterochannel electrical synapses in PLM. (A, B, C, E, H) Each point in the violin plot indicates number of innexin puncta observed in single PLM neurons. n > 20 neurons per group, unless specified. Statistical comparison between different groups were performed using the Fisher-Freeman-Halton exact test. n.s., non-significant, *p < 0.05, **p < 0.01, ***p < 0.001, ****p < 0.0001. Scalebar = 5 μm. (A) Punctate localization of endogenous INX-1::TAGRFP in *inx-1(amz01[inx-1::tagrfp])*, within PLM zone-1 electrical synapses (yellow arrowheads) remained intact in *unc-7(e5)* or *unc-9(e101)* single mutant and *unc-7(e5), unc-9(fc16)* double mutant animals. PLM axons were marked in blue. Quantifications are shown on the right. (B) Punctate localization of UNC-7::TAGRFP, in *unc-7(ot895[unc-7::tagrfp]) animals,* within PLM zone-1 electrical synapses (yellow arrowheads) remained intact in *inx-1(amz11)* animals, but were disrupted in *unc-9(e101)* animals. PLM axons were marked in blue. Quantifications are shown on the right. (C) Punctate localization of UNC-9::7xGFP, in *unc-9::7Xsplit-GFP animals,* within PLM zone-1 electrical synapses (yellow arrowheads) remained intact in *inx-1(amz11)* animals, but were disrupted in *unc-7(e5)* animals. PLM axons were marked by BFP expression. Quantifications are shown on the right. (D) Schematics showing domain structure of INX-1 and UNC-9 proteins, where EC1 and EC2 marks both the extracellular loops. In chimeric INX-1(\**UNC-9 EC*) protein, INX-1 EC1 and EC2 were replaced by corresponding UNC-9 extracellular loops. (E) PLM-specific expression of GFP::UNC-9 (*amzEx76*) in *unc-9(e101)* and INX-1(*UNC-9 EC)::GFP (*amzEx129*) in *unc-9(e101), inx-1(amz11)* animals were sufficient to rescue UNC-7::TAGRFP puncta in PLM synaptic zone-1. Quantifications are shown on the right. (F) Schematic of the *C. elegans* X chromosome showing positions of Floxed *unc-7(amz14 ot895[loxP::unc-7::tagrfp::loxP])*, FRT-flanked *inx-1(amz18 [FRT::inx-1::FRT])*, and FRT*-flanked *unc-9(amz32 [FRT*::unc-9::FRT*])* alleles. (G) UNC-7::TAGRFP puncta in the PLM synaptic zone-1 remained unaffected by Cre-recombinase mediated PLM-specific deletion of *unc-7* in unc-7 *(amz14 ot895); amzIs01[mec-18p::Cre::t2a::ebfp2]* animals. However, UNC-7::TAGRFP puncta were significantly affected upon Cre- and flippase-recombinase mediated simultaneous deletion of *unc-7, inx-1* and *unc-*9 specifically from PLM in unc-7 *(amz14 ot895) inx-1(amz18) unc-9(amz32)* animals. LoxP-or FRT-flanked alleles of *unc-7, inx-1* and *unc-9,* and PLM-specific Cre or flippase-recombinase expressing lines were validated using PCR (Figure S2E). (H) Quantification of the data shown in panel K. (I) Schematic showing co-immunoprecipitation of UNC-7 interacting innexins using anti-RFP nanobody in *unc-7(ot895)* animals. (J) Co-IP of UNC-7 as bait in *unc-7*(*ot895*), followed by mass-spectrometry identified UNC-9 but not INX-1 as UNC-7 interacting innexin. UNC-7 data used as positive control. Each data point represents the relative abundance normalized to total protein in each biological replicate. Pan-neuronally expressed TAGRFP in *amzIs09(UPNp::tagrfp*) was used as the negative control.

Next, to understand how UNC-7 participates in channel formation, we looked at the localization of endogenously tagged UNC-7 protein and found that it remained unaffected in *inx-1* mutant (Figure 2B). However, in *unc-9* mutants, the punctate localisation of PLM-synaptic UNC-7 was disrupted (Figure 2B). Similarly, UNC-9 localization remained unaffected in *inx-1* mutants, but its localization was disrupted in *unc-7* mutants (Figure 2E, F). These results suggested that UNC-7 and UNC-9 are mutually dependent on each other to form gap junction channels in PLM. Previous findings have shown that UNC-7 interacts with UNC-9 trans-synaptically to form heterotypic gap junction channels in other circuits^33,34,36^. To test whether UNC-7 and UNC-9 interact in trans to form heterotypic channels, we cell-specifically rescued UNC-9 in PLM and found that to be sufficient to rescue UNC-7 synaptic localisation in *unc-9* mutant animals, suggesting heterotypic interactions between them (Figure 2E). To further test whether UNC-7 and UNC-9 interact in a cis-manner to form heteromeric hemichannels, we used AlphaFold 3 modeling^38^ to analyse the putative structural stability of possible hexameric hemichannels formed by different combinations of INX-1, UNC-7 and UNC-9 (Figure S2C, D). Our modelling results suggested that it is more likely for INX-1, UNC-7 and UNC-9 to exist as independent hemichannels *in vivo* (Figure S2C, D). Further, previous reports have suggested that the extracellular loops of innexins/connexins are mediators of their docking specificity^39,40^. Accordingly, we replaced the two extracellular loops of INX-1 with corresponding sequences of UNC-9, to engineer a chimeric protein that resembled INX-1, but had docking capabilities of UNC-9, INX-1(*UNC-9EC). (Figure 2D). Cell-specific expression of this chimeric INX-1(*UNC-9EC) should rescue UNC-7 localization only if UNC-7 and UNC-9 interact in trans to form heterotypic channels. We found that PLM-specific expression of INX-1(*UNC-9EC) chimeric protein was sufficient to rescue UNC-7 localisation, suggesting UNC-7 and UNC-9 form heterotypic gap junction channels in PLM (Figure 2E).

Since UNC-7 and UNC-9 are present from both the synaptic partner sides, we hypothesized that PLM-electrical synapses may contain clusters of heterotypic UNC-7-UNC-9 channels that are reciprocal to one another. To test this hypothesis, we deleted *unc-7* specifically from PLM neurons using a loxP recombination-site-flanked conditional knockout allele of *unc-7* (*unc-7(amz14, ot895[loxP::unc-7::tagRFP-T::loxP]*) (Figure 2F, 2SE) and PLM-specific expression of Cre recombinase. This PLM-specific loss of *unc-*7 did not abrogate UNC-7 localization in PLM synapses. However, PLM-specific deletion of *unc-*7, *unc-9* and *inx-1* together, using loxP- or FRT-flanked alleles of *unc-7, inx-1* and *unc-9,* and PLM-specific simultaneous expression of Cre and flippase-recombinase, resulted in loss of UNC-7 puncta in PLM-synapse (Figure 2F-H, 2SE). Together, these results suggested that PLM-synapses are composed of reciprocally arranged heterotypic UNC-7-UNC-9 channels where UNC-7 and UNC-9 are present from both synaptic sides, along with independent INX-1 homotypic channels. As an independent line of evidence, we tested the interaction between UNC-7, UNC-9 and INX-1, via co-immunoprecipitation of endogenously tagged-UNC-7, *unc-7(ot895[unc-7::tagRFP-T::loxP]*), followed by mass spectrometric analysis (Figure 2I). Our biochemical data suggested that while UNC-7 strongly interacts with UNC-9, it does not interact with INX-1 (Figure 2J). Together, our data suggest that PLM neurons form novel “Heterochannel Electrical Synapses”, where individual electrical synapses can be formed by clustering molecularly distinct gap junction channels.

### Distinct channels in a synapse function redundantly to elicit touch response

Having defined the molecular composition of heterochannel PLM-synapses, we wanted to understand how individual channel-types in heterochannel-synapses contribute to overall synaptic function, to ultimately understand the role of PLM-electrical synapses in posterior mechanosensation. Based on the molecular composition of heterochannel PLM-synapses, we hypothesized that individual innexin mutants could be insufficient to abrogate PLM-electrical synaptic function due to functional compensation from the other channel types. To test this hypothesis, we generated pairwise double mutants of innexins and assayed them for touch sensitivity (Figure 3A). Aligning with our previous observations, animals mutant for both *unc-7* and *unc-9* did not show any significant defects in posterior touch response, similar to individual mutants (Figure 3A). These results further suggested that UNC-7 and UNC-9 function as single-channel units. However, consistent with the heterochannel synapse model, double mutants of *inx-1* with either *unc-7* or *unc-9*, where all the PLM gap junction channel-types were disrupted, showed almost complete loss of posterior mechanosensation (Figure 3A). Since all three innexins under consideration are widely expressed in the nervous system, we wanted to ensure that the defect in touch sensitivity is arising specifically out of TRN electrical synapse perturbation. To achieve this, we deleted either *unc-7* alone or *inx-1* and *unc-9* together specifically in TRNs using conditional knockout alleles of *unc-7, inx-1* and *unc-9* (Figure 2F), as previously mentioned. Resembling whole animal mutant for *unc-7*, *unc-7(e5)*, PLM-specific knockout of *unc-7* also did not affect posterior mechanosensation (Figure 3A). However, similar to *unc-9(e101) inx-1(amz11)* double mutant animals, PLM-specific knockout or *unc-9* and *inx-1* together showed a significant reduction in posterior mechanosensation (Figure 3A). In a complementary experiment, we found that PLM-specific rescue of *unc-9* in *unc-9(e101) inx-1(amz11)* double mutant animals resulted in significant improvement in the touch response behaviour (Figure S3A). These results together showed that PLM-electrical synapses drive the posterior low-threshold mechanosensation, as originally hypothesized^20^. Furthermore, our results suggested that molecularly distinct channel-types in a heterochannel electrical synapse can be functionally redundant.

**Figure 3.**
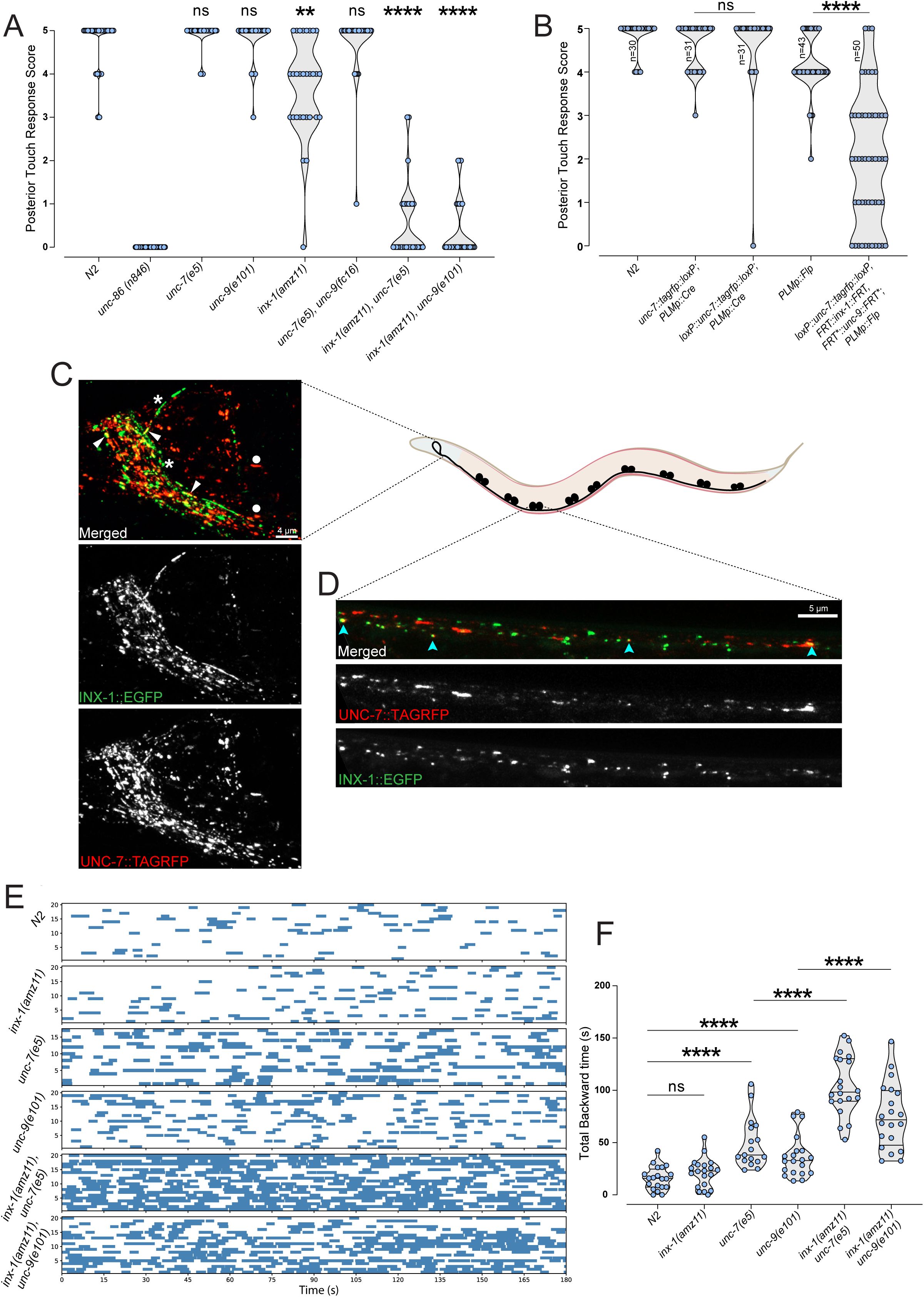
*inx-1, unc-7* and *unc-9* function redundantly in the gentle touch and locomotion circuits. (A-B) Posterior low-threshold touch response in young adult animals. Each data point represents response of a single animal. Statistical comparison between different groups were performed using the Fisher-Freeman-Halton exact test. n.s., non-significant, *p < 0.05, **p < 0.01, ***p < 0.001, ****p < 0.0001. (A) *inx-1(amz11), unc-7(e5) and inx-1(amz11), unc-9(e101)* double mutant animals showed significant enhancement of the posterior touch response defect compared to *inx-1(amz11)* single mutant animals, similar to the *unc-86(n846)* positive control animals. *unc-7(e5)*, *unc-9(e101)* single mutant or *unc-7(e5), unc-9(e101)* double mutant animals showed non-significant affect. Single mutant animal data are same as shown in Figure 1A. n=30 for each genotype. Data for single mutants were repurposed from Figure 1A. (B) Flippase-recombinase mediated PLM-specific deletion of *inx-1* and *unc-9* together significantly affected the posterior touch response in *inx-1(amz18),unc-9(amz32); amzEx79[mec-4p::Flp::t2a::ebfp2]* animals. Cre-recombinase mediated PLM-specific deletion of *unc-7* in *unc-7(amz14 ot895); amzIs01[mec-18p::cre::t2a::ebfp2]* animals had no significant effect on posterior touch response. (C) Representative punctate expression of endogenously-tagged INX-1::GFP and UNC-7::TAGRFP in the head of an adult animal, *inx-1(amz21[inx-1::gfp]), unc-7*(*ot895[unc-7::tagrfp]*). White arrowheads indicate colocalization of INX-1 and UNC-7 within representative puncta. Asterisks indicate puncta having only INX-1 and solid circles indicate puncta having only UNC-7. Image obtained using Airyscan mode with 63X oil-immersion objective. Scalebar = 4[μm. (D) Representative image showing co-localization of UNC-7 and INX-1 puncta in the ventral nerve cord (cyan arrowheads), in *unc-7(ot895), inx-1(amz21)* animals. Scalebar = 5 μm. (E) Locomotion assay using the WormLab automated tracking system (see Methods). Each row in the raster plot represents the trajectory of a single young adult animal. Blue bars denote time spent in backward locomotion during a 3-minute recording period. *inx-1(amz11), unc-7(e5) and inx-1(amz11), unc-9(e101)* double mutant animals showed significantly enhanced backward locomotion, compared to N2 or respective single mutant animals. (F) Quantification of total time spent in backward locomotion. Each circle represents the experimental mean of a single animal. Data were tested for normality using the Shapiro–Wilk test. Statistical comparisons were performed using Welch’s *t*-test (for normally distributed samples) or the two-tailed Mann–Whitney *U* test (for non-normally distributed samples). *inx-1(amz11), unc-7(e5) and inx-1(amz11), unc-9(e101)* double mutant animals showed significantly enhanced backward locomotion, compared to respective single mutant animals. n>17 for each genotype.

As *inx-1, unc-7* and *unc-9* are all widely expressed in the nervous system, our observations in the PLM-circuit prompted us to ask whether these innexins also cooperate outside the TRN-circuit by potentially forming similar heterochannel synapses, to regulate other behaviours in *C. elegans.* To address this question, we looked at the co-localisation between these innexins across the nervous system. Previously, using transgenic reporter lines, UNC-7 and UNC-9 have been shown to extensively colocalise in the *C. elegans* nervous system^33^. Using our endogenously tagged alleles of UNC-7 and INX-1, we examined their colocalization in the nerve ring region of the animals. We observed that while in most cases UNC-7 and INX-1 formed spatially separate puncta, there exists sporadic colocalization between these innexins in specific synapses (Figure 3C, Movie S1), suggesting synapse-specific collaborative function between them. Previously, UNC-7 and UNC-9 were shown to extensively colocalize in the ventral nerve cord (VNC) to form heterotypic rectifying channels that are necessary for proper locomotion^33,34,36^. Using endogenously tagged reporter alleles, we found that INX-1 and UNC-7 also stereotypically colocalize in some electrical synapses present along the VNC (Figure 3D).

To test whether INX-1 could also function redundantly with UNC-7 or UNC-9 outside of the mechanosensory circuit, we quantitatively assessed the locomotory behaviour of freely moving young adult animals that are mutant for either single or double mutants for *inx-1*, *unc-9* and *unc-7*, using an automated worm tracker, the Wormlab2 system. We observed that the *inx-1*, *unc-7* or *inx-1, unc-*9 double mutant animals showed significantly enhanced defects in multiple aspects of the worm locomotion, compared to *unc-7* or *unc-*9 mutant animals (Figure 3E, F, S3B-D). Especially, *inx-1*, *unc-7,* or *inx-1, unc-*9 double mutant animals spent significantly more time performing reversals compared to animals that are either single mutants for *inx-1, unc-7* and *unc-9,* or double mutants for *unc-7*, *unc-9* (Figure 3E, F). These results suggest that similar to TRN synapses, this aggravation of uncoordinated phenotype could arise from the disruption of putative heterochannel synapses present in the locomotion circuit.

### Molecularly distinct channels within an heterochannel electrical synapse are transported via distinct mechanisms

Having established the functionally important heterochannel synaptic composition of PLM-electrical synapses, we next sought out to understand the molecular mechanisms regulating formation of these synapses. We first focused on the transport mechanisms of molecularly distinct gap junction channels to heterochannel PLM-synapses. Unlike transport of chemical synaptic components, including synaptic vesicles, neurotransmitter receptors, transport mechanisms for innexins remain much understudied. Transport of UNC-9 in PLM axons has been shown to be independent of kinesin-3 family member, UNC-104/KIF1A, and kinesin-1 family member, UNC-116^22^; both of which regulate trafficking of synaptic vesicles and mitochondria in PLM axons, respectively ^41–43^. To systematically determine how INX-1 and UNC-7/UNC-9 channels get transported to the heterochannel PLM-synapses, we focused specifically on the kinesin-family of motor proteins that are primary regulators of axonal anterograde vesicular cargo transport. Using RNA-interference (RNAi), we systematically knocked down 19 out of 21 kinesin-family of proteins encoded in the *C. elegans* genome^44,45^ and analysed the effects on the localization of INX-1::TagRFP and GFP::UNC-9 into electrical synapses within PLM synaptic zone. We found that knocking down of three distinct kinesins: a kinesin-3 family member, *klp-4*/KIF13, KIF14 and two of the kinesin-14 family members, *klp-3/KIFC2, KIFC3* and *klp-15/KAR3* independently affected punctate localizations of INX-1::TAGRFP-T in PLM synapses, where significant number of PLM axons showed either complete loss or reduced number of INX-1 puncta in synaptic zone-1 and 2 (Figure 4A; S4A,B). Consistent with our RNAi-data, we observed either complete loss or reduced localization of endogenously tagged INX-1::TAGRFP-T in PLM synaptic zones in *klp-4(ok3537*) mutant animals [zone 1: 85.2% and zone 2: 73.3% of neurons showing defect] (Figure 4B,C) and *klp-4(tm2114)* [zone 1: 51.6% of neurons showing defect] (Figure S4C, D). Similarly, *klp-3(ve638*) mutants also showed either reduced localization or complete loss of endogenous INX-1::TAGRFP-T [zone 1: 51.9% and zone 2: 54.5% of neurons showing defect] (Figure 4B,C). On the other hand, *klp-15(ok1958)* mutants did not show a significant loss of endogenous INX-1::TAGRFP-T localisation (discussed later). These results suggested that two kinesins belonging to two distinct conserved families regulate the trafficking of INX-1 to electrical synapses in PLM axons.

**Figure 4.**
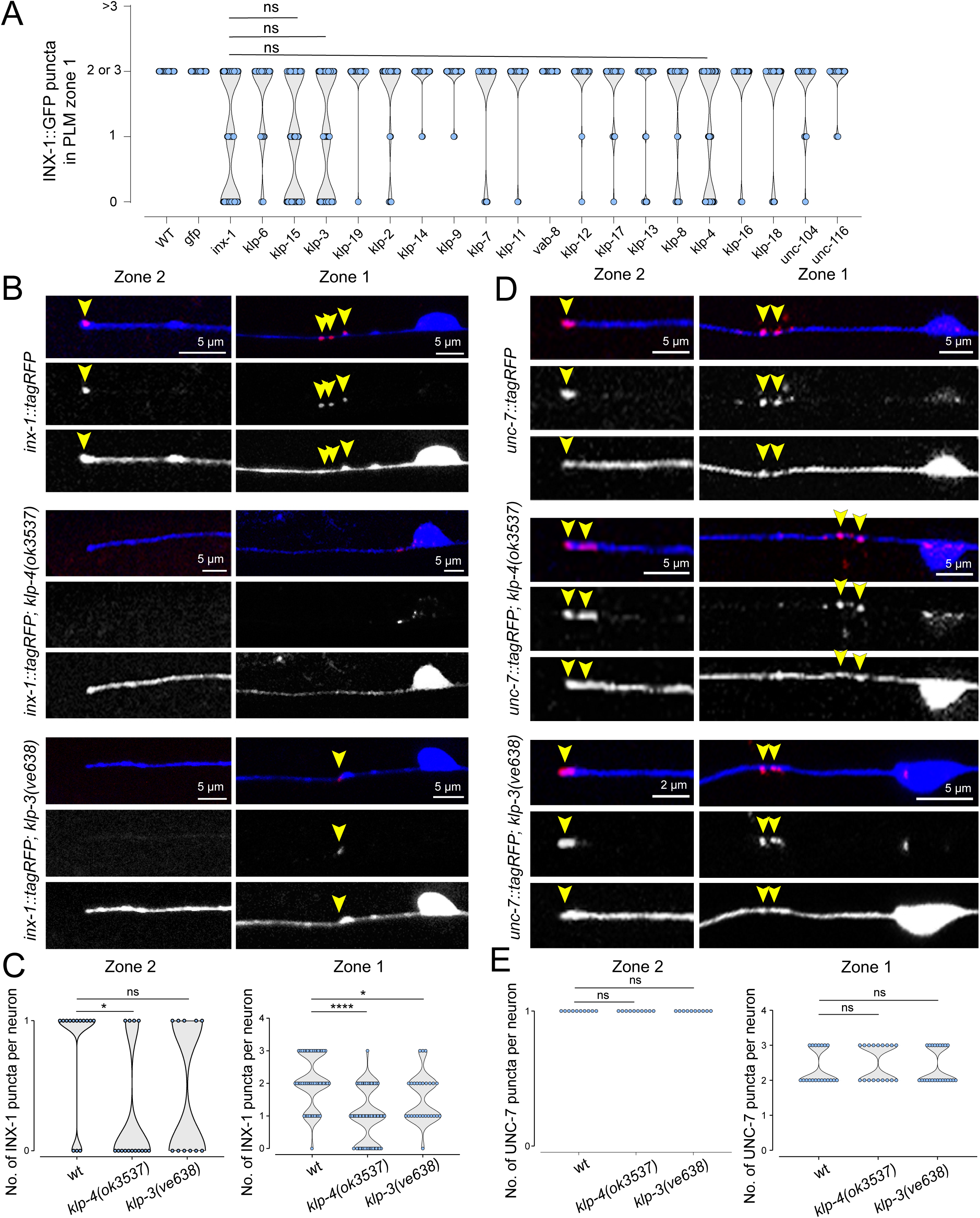
Synaptic localization of *inx-1* and *unc-7* depend on distinct motor proteins. (A) Each data point represents number of distinct INX-1::TagRFP (*amz66*) puncta observed within synaptic zone 1 of a single PLM neuron. Statistical comparison between different groups were performed using the Fisher-Freeman-Halton exact test. n.s., non-significant, *p < 0.05, **p < 0.01, ***p < 0.001, ****p < 0.0001. PLM neuron-specific RNAi-mediated knockdown of *inx-1* in *sid-1(pk3321) him-5(e1490) V; lin-15B(n744) X; uIs71 [(pCFJ90) myo-2p::mCherry + mec-18p::sid-1]; amzEx66[mec-4p::inx-1a::tagrfp; mec-4p::unc-1::gfp; ttx-3p::gfp]* served as the positive control, while RNAi against *gfp* served as the negative control. Knockdown of kinesin genes, *klp-15, klp-3* and *klp-4* resulted in similar reduction in INX-1::TagRFP puncta, comparable to the positive control. n>30 for all knock-downs. (B) Punctate localization of endogenous INX-1::TAGRFP (yellow arrowheads) in *inx-1(amz08[inx-1::tagrfp])*, within PLM synaptic zone-1 and -2 were significantly reduced in *klp-4(ok3537)* and *klp-3(ve338)* mutant animals. PLM axons were marked in blue. (C) Quantification of the data shown in panel A. (D) Punctate localization of endogenous <u>UNC</u>-7::TAGRFP (yellow arrowheads) in *unc-7(ot895[unc-7::tagrfp])*, within PLM synaptic zone-1 and -2 remained unaffected in *klp-4(ok3537)* and *klp-3(ve338)* mutant animals. PLM axons were marked in blue. (E) Quantification of the data shown in panel D.

Intriguingly, mutations in *klp-4* and *klp-3* did not significantly affect synaptic localization of UNC-7 and UNC-9 in PLM neurons (Figure 4D, E, S4E, F), nor did knockdown of other kinesins using RNAi (Figure S4F). Taken together, these results suggested that INX-1 and UNC-7-UNC-9 gap junction channels depend on distinct molecular motors for localization to heterochannel electrical synapses in PLM.

### UNC-7/UNC-9 and INX-1 channels in PLM heterochannel synapses are regulated by distinct mechanisms

The heterochannel nature of PLM electrical synapses and their distinct mode of synaptic transport prompted us to ask how these molecularly distinct channels are being regulated by other known mechanisms of electrical synapse assembly-disassembly. Previous reports suggested that UNC-9 localization in PLM synapses is regulated by unc-44/Ankyrin, UNC-33/CRMP and VAB-8/Atypical kinesin-pathway, which according to the proposed model, is required for the turnover of UNC-9 channels from PLM synapses^22^. As a result, mutations in unc-44/UNC-33/VAB-8 pathway genes results in supernumerary UNC-9 puncta phenotype in the PLM synaptic zones^22^. Since UNC-7 and INX-1 also localise to the same synaptic puncta, we asked whether the unc-44/UNC-33/VAB-8 pathway also similarly regulates UNC-7 and INX-1 dynamics in PLM-synapses. Our results showed that endogenously-tagged UNC-7 also forms supernumerary puncta at PLM synaptic zones, when *unc-44* or *unc-33* or *vab-8* were mutated, reminiscent of UNC-9 defects (Figure 5A-D) and consistent with our previous results suggesting UNC-7 and UNC-9 work together in PLM-synapses. Surprisingly, localisation of endogenously-tagged INX-1 remained largely unaffected in animals mutant for *unc-44* or *unc-33* or *vab-8* (Figure 5C-D). These findings confirmed that molecularly distinct gap junction channels within individual heterochannel synapses are differentially regulated by the potential mechanisms associated with channel turnover.

**Figure 5.**
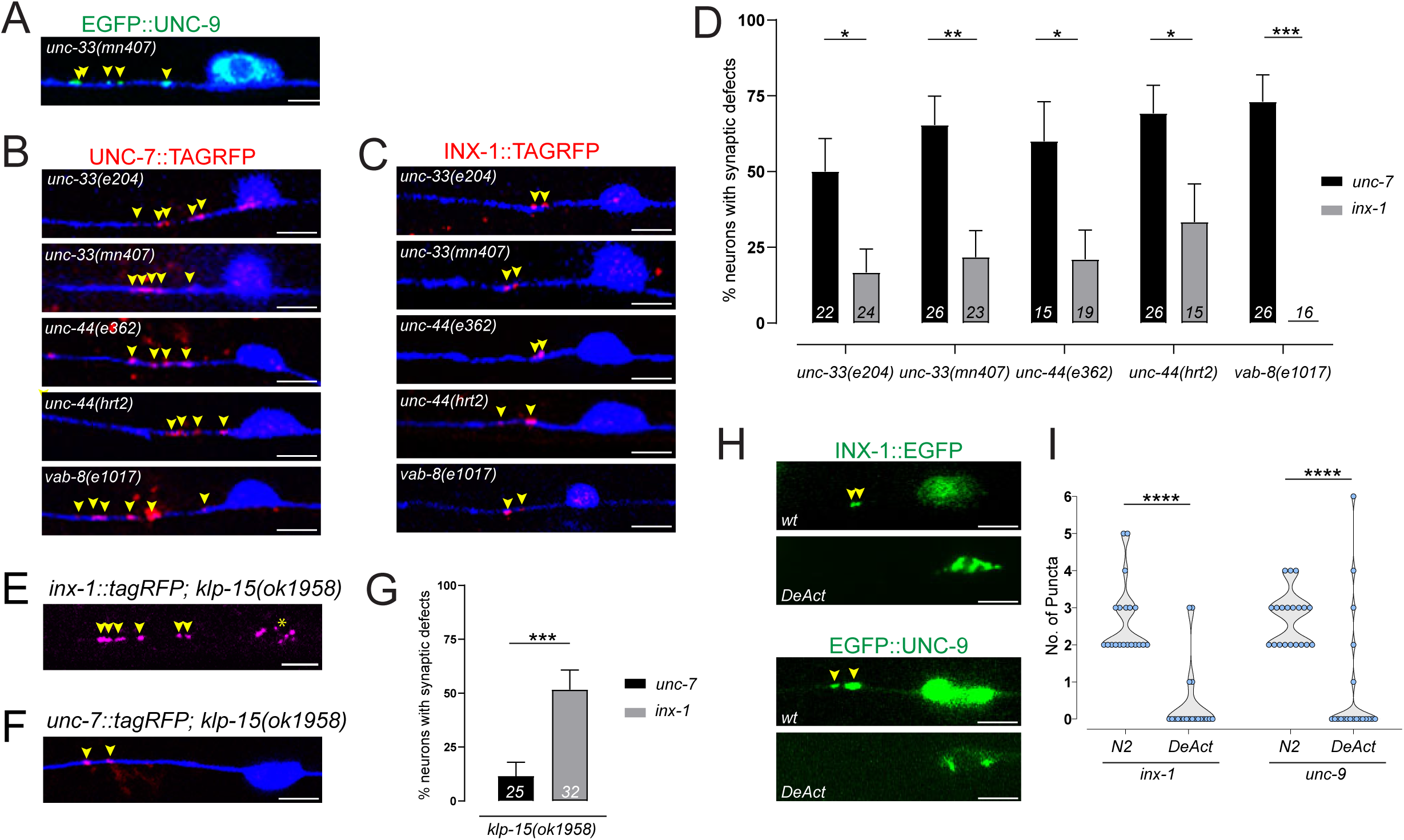
Number of molecularly distinct gap junction channels in PLM are restricted by distinct regulatory pathways. (A-D, G) PLM axons are marked in blue. Scale bar = 5 μm. (A) GFP::UNC-9, *amzEx76[mec-4p::GFP::3XGS::unc-9],* forms supernumerary puncta (yellow arrowheads) within PLM synaptic zone-1 in *unc-33(mn407)* animals, as previously reported^22^. (B) In *unc-33*, *unc-44*, and *vab-8* mutant animals, UNC-7, *unc-7(ot895[unc-7::tagRFP]),* also forms supernumerary puncta (yellow arrowheads) within PLM synaptic zone-1. (C) In *unc-33*, *unc-44*, and *vab-8* mutant animals, INX-1, *inx-1(amz01[inx-1::tagRFP]),* puncta (yellow arrowheads) within PLM synaptic zone-1 remained largely unaffected. (D) Quantification of neurons exhibiting supernumerary puncta defects of UNC-7::TAGRFP and INX-1::TAGRFP in *unc-33*, *unc-44*, and *vab-8* mutant backgrounds. *n* > 15 neurons per allele per gene. Error bars represent the standard error of the mean (SEM). Statistical tests were performed using the two-tailed Fisher’s exact test. n.s., non-significant, *p < 0.05, **p < 0.01, ***p < 0.001, ****p < 0.0001. (E) In *klp-15(ok1958)* mutant animals, INX-1, *inx-1(amz01[inx-1::tagRFP]),* forms supernumerary puncta (yellow arrowheads) within PLM synaptic zone-1. Yellow asterisk indicates cell-body accumulation of INX-1 protein. (F) In *klp-15(ok1958)* mutant animals, UNC-7, *unc-7(ot895[unc-7::tagRFP]),* puncta (yellow arrowheads) within PLM synaptic zone-1 remained unaffected. (G) Quantification of neurons exhibiting supernumerary puncta defects of UNC-7::TAGRFP and INX-1::TAGRFP in *klp-15(ok1958)* mutant backgrounds. Error bars represent the standard error of the mean (SEM). Statistical tests were performed using the two-tailed Fisher’s exact test. n.s., non-significant, *p < 0.05, **p < 0.01, ***p < 0.001, ****p < 0.0001. (H) In DeActs background, both INX-1, (*amzEx110*), and UNC-9 (*tbEx448*) fail to localise in the PLM synaptic Zone-1. (I) Quantification of neurons exhibiting loss of INX-1 and UNC-9 from PLM synaptic Zone-1 in DeActs background. Statistical tests were performed using the Fisher-Freeman-Halton exact test. n.s., non-significant, *p < 0.05, **p < 0.01, ***p < 0.001, ****p < 0.0001. (n=20 neurons)

During analysis of kinesin mutants, we found that endogenously-tagged INX-1 forms supernumerary puncta in PLM-neurons in *klp-15(ok1958)* mutant animals (in 53.12% of *klp-15* mutant PLM-neurons) (Figure 4E, G), reminiscent of the UNC-7 and UNC-9 expression defect in *unc-44*/*unc-33*/*vab-8-*mutant animals. However, *klp-15(ok1958)* mutation did not affect the localization of endogenously-tagged UNC-7 in PLM synaptic zones (Figure 4F, G). These data together suggest that two independent mechanisms regulate the number of molecularly distinct gap junction puncta within individual heterochannel electrical synapses in PLM.

The junctional maintenance gap junctions require structural support from the F-actin mediated cytoskeletal network, and other actin-associated scaffolding proteins^46^. Given above mentioned distinctions in the associated machinery with distinct innexins, we asked whether the disruption of cytoskeletal assemblies has a differential impact on these innexins. Towards this end, we exploited DeAct, a genetically encoded polypeptide that disrupts the F-actin network^47^. We observed that punctate localisation of both UNC-9 and INX-1 to PLM synapses were disrupted in thd DeActs background (Figure-5H, I), suggesting generic role of the F-actin network in the assembly of heterochannel synapses. A recent study showed that and the assembly of UNC-9/UNC-7 heterotypic electrical synapses in *C. elegans* motor circuit is regulated by cAMP-levels, maintained by cAMP-antagonist, phosphodiesterase *pde-1*^16^. Our results suggested that UNC-7 and INX-1 localizations in PLM-synapses are independent of *pde-4(ce268)* mutants, the most abundantly expressed phosphodiesterase in PLM neurons^12^ (Figure S5B, C).

Taken together, these results suggested that the dynamics of INX-1 and UNC-7/UNC-9 channels present within individual heterochannel PLM-synapses depend on specific, but distinct molecular mechanisms.

### Molecular composition of PLM-BDU heterochannel synapse gets reversibly altered in dauer-stage animals

Connexins and innexins have been known to be expressed in a temporally regulated manner^48,49^. Additionally, our previous work showed that the nervous system wide innexin expression atlas undergoes substantial alteration in a neuron-specific manner upon entry into the dauer diapause stage, under harsh environmental conditions^13^. These alterations in innexin expression ultimately dictate electrical synaptic connectivity as well as specific aspects of dauer behaviour^13^. These observations prompted us to ask whether the molecular composition of heterochannel electrical synapses in PLM can also undergo alteration at the dauer stage, under adverse environmental conditions. We found that INX-1 and UNC-7 continue to colocalize at the PLM-synapses present within the synaptic zone 1 also in dauer animals (Figure 6A). However, contrastingly, INX-1 localization appeared selectively downregulated in synaptic zone 2 in PLM-BDU synapses, while UNC-7 continued to be localized there (Figure 6B, C). Furthermore, this dauer-specific down-regulation of INX-1 localization at the PLM-BDU synapse was found to be reversible and rebounded within 12 hours of feeding, when dauer animals resumed development under favourable conditions (Figure 6C). Overall, these results suggested that the molecular composition of heterochannel synapses are dynamic and reversible in a synapse-specific manner even within individual neurons.

**Figure 6.**
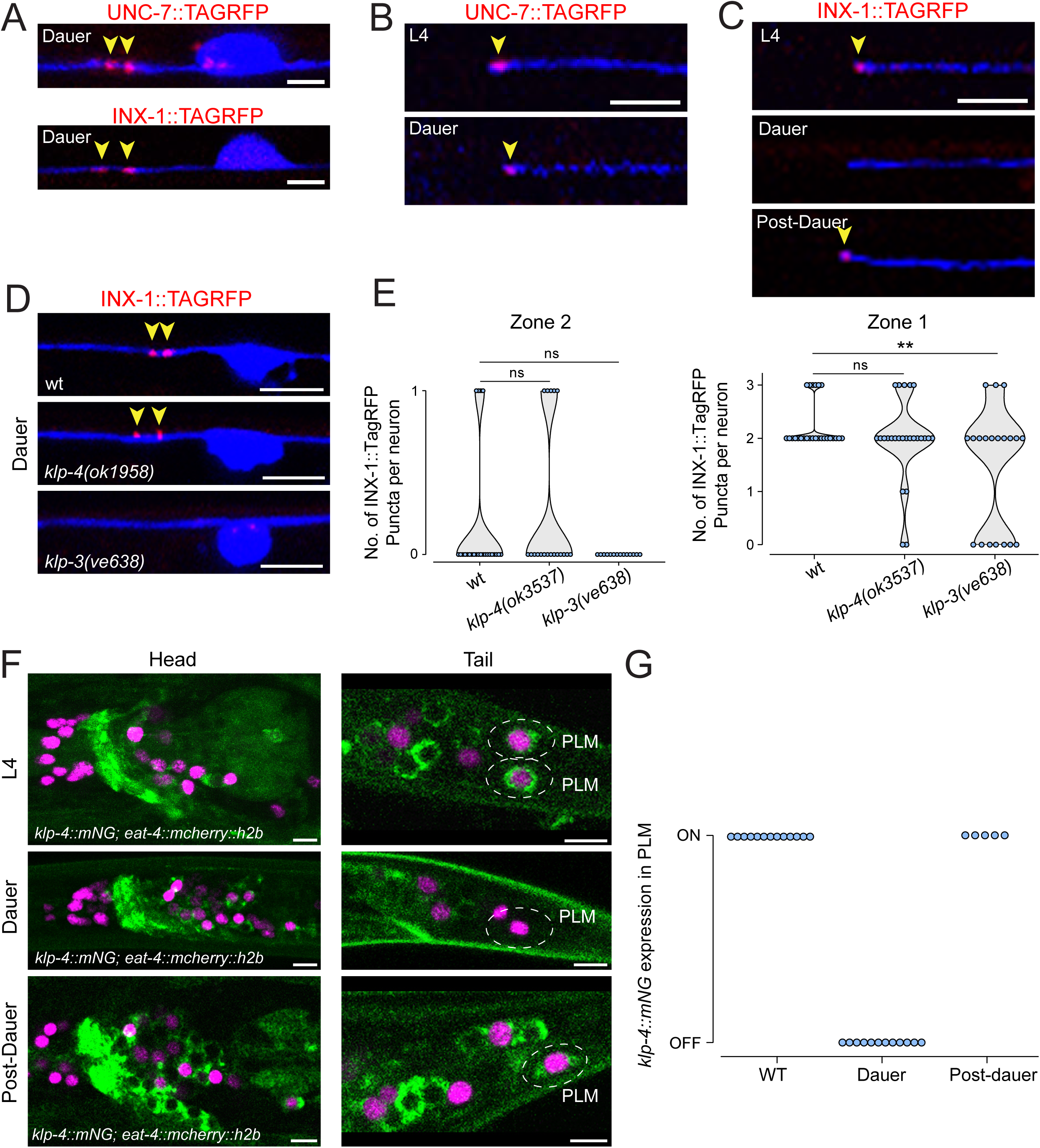
Number of molecularly distinct gap junction channels in PLM are restricted by distinct regulatory pathways. (A-D) PLM axons are marked in blue. Scale bar = 5 μm. (A) UNC-7, *unc-7(ot895[unc-7::tagRFP]),* and INX-1, *inx-1(amz01[inx-1::tagRFP]),* puncta (yellow arrowheads) within PLM synaptic zone-1 remained unaltered in dauer animals. (B) UNC-7, *unc-7(ot895[unc-7::tagRFP]),* puncta (yellow arrowheads) within PLM synaptic zone-2 remained unaltered in dauer animals. (C) INX-1, *inx-1(amz01[inx-1::tagRFP]),* puncta (yellow arrowheads) within PLM synaptic zone-2 gets selectively downregulated in dauer stage, while it is restored in post-dauer animals. (D) In the dauer stage, localization INX-1::TAGRFP (yellow arrowheads), *inx-1(amz08[inx-1::tagrfp])*, puncta within PLM synaptic zone-1 were not affected in *klp-4(ok3537)* mutant animals, but were affected in *klp-3(ve338)*. (E) Quantification of punctate INX-1::TAGRFP localization within PLM synaptic zones in *klp-4(ok3537)* and *klp-3(ve338)* mutant animals. n>15 neurons. (F) Expression of endogenously-mNeongreen tagged KLP-4, (*klp-4(cp303[klp-4::mNG]),* gets selectively downregulated in PLM neurons in dauer, while continue to be expressed broadly in the nerve ring. KLP-4::mNG expression in PLM gets restored in post-dauer animals. PLM were identified using co-expression of *eat-4* reporter^78^. (G) Quantification of the data shown in panel F.

Since we identified that kinesin motors distinctly regulate trafficking of INX-1 and UNC-7/UNC-9 channels to PLM synapses in non-dauer stages, we further assessed the involvement of *klp-4* and *klp-3* on INX-1 and UNC-7/UNC-9 localization in dauer animals. We found that, unlike non-dauer animals, localization of endogenously-tagged INX-1 at PLM synapses within the zone-1 was completely unaffected in *klp-4(ok3537*) mutant dauer animals (Figure 6D, E), but continued to be affected in *klp-3(ve638*) mutant dauers [zone 1: 35% of neurons showing defect] (Figure 6D, E). Furthermore, to understand the basis of dauer-stage specific altered kinesin dependency of INX-1 trafficking, we looked at the expression of *klp-4* using an endogenously tagged allele*, klp-4(cp303[klp-4::mNG-C1^3xFlag])*^50^*. klp-4* is broadly expressed in head neurons and in several tail neurons, including PLM neurons (Figure 6F). However, our results showed that in dauer stage animals, *klp-4* expression gets selectively downregulated in tail neurons, including PLMs, while continue to be expressed broadly in the head neurons (Figure 6F, G). Moreover, dauer stage-specific down-regulation of *klp-4* expression in PLM neurons was reversible and regained within 12 hours of feeding, when dauer animals recovered under favourable conditions (Figure6F, G); reminiscent of the reversible nature of INX-1 localization in PLM-BDU synapse in post-dauer stage (Figure 6C). Altogether, our results demonstrate that expression of specific kinesin-motor proteins undergoes neuron-type specific modulation under an altered environmental condition, which potentially could be responsible for the altered molecular composition of heterochannel synapses in distinct electrical synapses in specific neuron-types.

### Cx35b and Cx43 colocalize in electrical synaptic puncta in the Purkinje neurons of developing zebrafish larvae

Similar to *C. elegans* neurons, previous reports suggested that individual neuron classes in vertebrate model systems can also express multiple connexin genes across developmental stages^15,51,52^. Our study identifying functionally important heterochannel synapses in the *C. elegans* nervous system prompted us to examine the conservation of this electrical synaptic organization principle in vertebrate neurons, where multiple connexins are co-expressed. Previous studies using zebrafish larvae as a model to study combinatorial connexin expression code in distinct neuron types^15,51^, showed that over 10 connexins are expressed in the zebrafish brain^51^. In the larval zebrafish, Purkinje neurons have been previously shown to form electrical synapses composed of Cx35b (a homolog of human Cx36) that are important for dendritic arborization and chemical synapse formation^53^ (Figure 7A). We referred to the single cell transcriptomic data from the zebrafish cerebellum^14^, and found that indeed there are multiple connexins expressed in Purkinje neurons (Figure 7B), making them a suitable candidate to study interactions between different connexins. Upon analyzing the RNA sequencing data from Takeuchi et al.^14^, we found that Cx43 and Cx43.4 are the most heavily expressed connexins in these neurons, prompting us to ask if these connexins can interact with Cx35b to form putative heterochannel synapses. To this end, we examined the localization of Cx43 and Cx43.4 with respect to the previously characterized Cx35b. We sparsely expressed GFP-tagged Cx43 and Cx43.4 along with mCherry-tagged Cx35b in a pair-wise manner using the Purkinje-specific, Ca8-cfos promoter^25^ and looked at their localization in 4-7day post fertilization (4-7 dpf) developing larvae. Similar to Cx35b, we found that both Cx43 and Cx43.4 localize in punctate structures along the dendrite as well as in the soma of Purkinje neurons (Figure 7C, D). We observed extensive co-localization of Cx43 with Cx35b in both soma and dendrites of Purkinje neurons (Figure 7C). On the other hand, Cx43.4 rarely colocalized with Cx35b either in the soma or in the dendrite (Figure 7D). Moreover, a previous TurboID-based biochemical interactome study of Cx35b in zebrafish did not identify either Cx43 or Cx43.4 as Cx35b interactors^54^. We conclude that Cx43 and Cx35b could be part of putative heterochannel synapses in zebrafish Purkinje neurons, but are unlikely to form heteromeric channels. Therefore, heterochannel synapses may represent a conserved feature across evolution to diversify electrical synaptic transmission.

**Figure 7.**
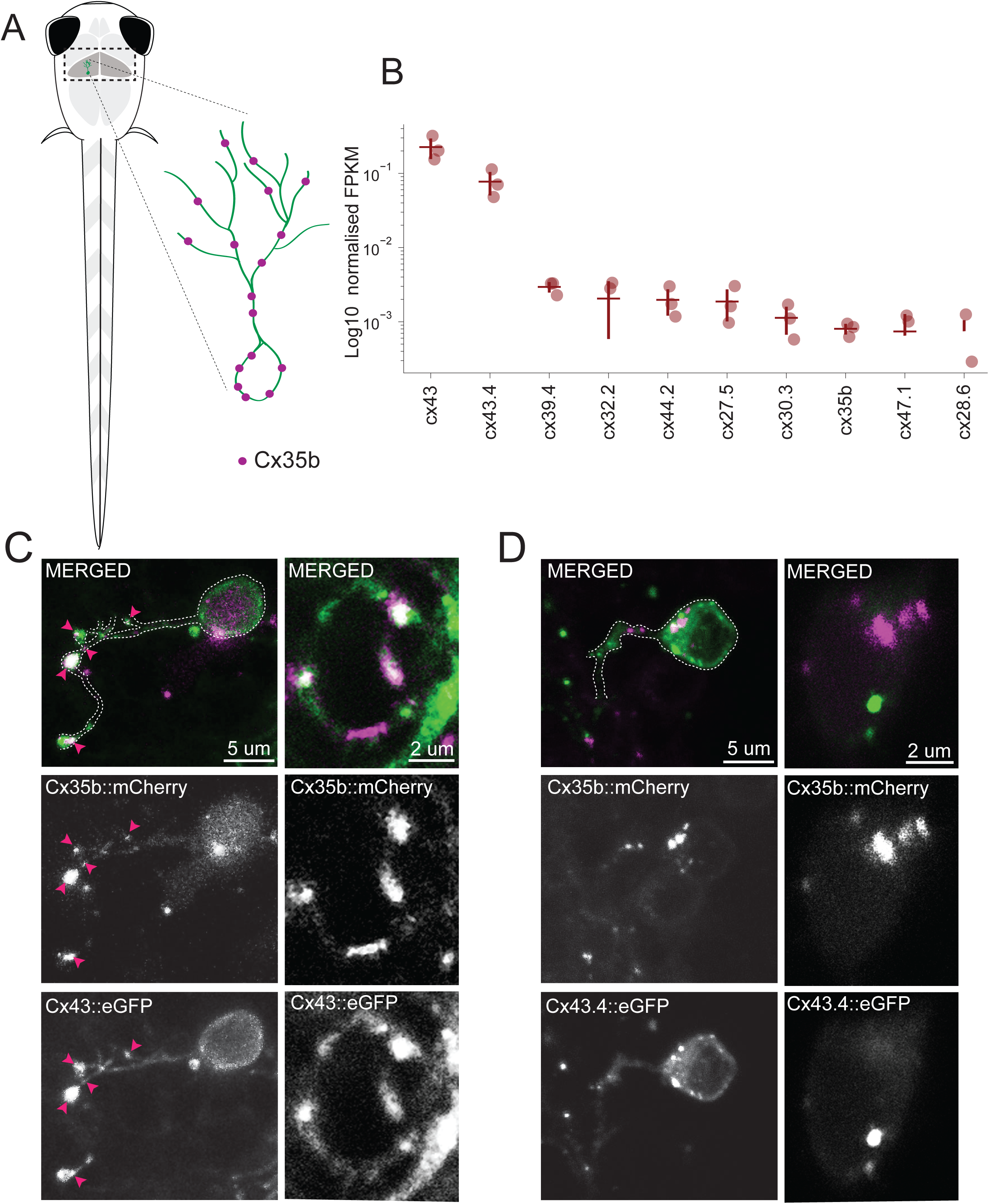
Cx35b and Cx43 co-localize at putative electrical synapses within zebrafish Purkinje neurons. (A) Schematic of zebrafish larva highlighting the anatomical position of cerebellar Purkinje neurons. The inset shows schematic of a single Purkinje neuron. Magenta dots represent putative Cx35b-labeled electrical synaptic puncta along the soma and dendrite. (B) Scatter plot showing log-normalized expression value of specific connexins in larval Purkinje neurons, obtained from single-cell RNA-sequencing dataset^14^. Each point represents a biological replicate. Normalization was done with GAPDH within Purkinje neurons. Horizontal bars indicate the mean; error bars denote SEM. (C) Representative confocal images showing co-localization of Cx35b::mCherry and Cx43::eGFP puncta along Purkinje neuron dendrites (magenta arrowheads) and soma. (D) Representative confocal images showing separate punctate expression of Cx35b::mCherry and Cx43.4::eGFP in the Purkinje neuron dendrites and soma.

## DISCUSSION

The fast-acting nature of the electrical synapses has been generally associated with the escape response in vertebrates and invertebrates. While previous cell-ablation and connectome studies in *C. elegans* suggested that the electrical synapses between touch-receptor neurons and forward pre-motor interneuron play key roles in guiding the low threshold mechanical stimulation mediated forward escape behaviour ^20^, genetic loss of function studies of innexins showed rather mild or no phenotypes^23,24^. Through our studies in the mechanosensory circuit in *C. elegans*, we have uncovered a novel configuration of electrical synapses wherein individual synapses are formed by clustering molecularly distinct gap junction channel-types, which we term heterochannel electrical synapses. We have also shown that the gap junction channel code of the heterochannel electrical synapses plays crucial roles in mediating the posterior touch-response behaviour, explaining the previously mentioned discrepancies. We also provided evidence as well as functional correlates of heterochannel electrical synapses in the locomotory circuits in *C. elegans*. In line with our findings, a previous study also reported enhanced uncoordinated locomotion in either *inx-1*, *unc-7* or *inx-1, unc-*9 double mutant animals compared to the single mutant animals in the RIM-AVA/AVE circuit that controls reversals^55^. Furthermore, investigating PLM electrical synapses, we have also shown that the heterochannel synaptic composition can undergo dynamic modulation in response to changes in environmental conditions. Extending beyond the *C. elegans* nervous system, we also provide evidence for putative heterochannel electrical synapses in Purkinje cells in larval zebrafish cerebellum.

Our work provides the basis to understand several critical features of the electrical connectome. As individual neurons often express multiple innexin or connexin genes and their splice isoforms^12,13,25^, this work provides a framework to understand how neurons may utilize their innexin or connexin repertoire for functional robustness and diversity. As the molecular composition of gap junction channels has been shown to dictate their functional properties^37,56,57^, we hypothesize that the combinatorial gap junction code of heterochannel electrical synapses may allow further diversification of ways in which signals can flow through the synapse, and from a broader viewpoint, through the electrical connectome. This synapse-specific combinatorial gap junction code, along with the neuron-specific combinatorial innexin expression code, may ultimately determine the synaptic partner choices and functional properties.

Our finding also opens up the possibility of potential modulation of the gap junction channel code of individual electrical synapses. Co-localization of distinct types of neurotransmitter receptors and their stoichiometric alteration has been well appreciated in the case of chemical synapses, especially co-localization and alteration of AMPA and NMDA receptors in glutamatergic synapses have been proven to be important contributors of functional diversity^58,59^. To gain a deeper understanding of the cell biological processes regulating the formation of heterochannel synapses, we analysed the synaptic transport mechanisms of channels formed by INX-1 and UNC-7/UNC-9 within PLM synapses, a process that has not been fully understood. Our findings identified that INX-1 channels are transported by two conserved kinesin motors, a kinesin-3 family protein, KLP-4, and a kinesin-14 family member, KLP-3, in a neuron-specific manner. Intriguingly, trafficking of other gap junction channels formed by UNC-7 and UNC-9 to the same synapse is independent of KLP-4 and KLP-3. KLP-4 is a microtubule plus-end directed anterograde motor^42,60^, which has also been shown to regulate anterograde trafficking of glutamate receptor, GLR-1 in *C. elegans* ventral nerve cord neurons^61^. On the other hand, members of the kinesin-14 family play major roles in spindle assembly and primarily regulate microtubule minus-end directed retrograde movement^62–64^. However, kinesin-14 family members have also been shown to promote context-dependent anterograde cargo movement^65,66^. This suggests that two kinesins belonging to two distinct conserved families regulate trafficking of specific types of innexin channels to heterochannel electrical synapses. On the other hand, it was shown that cytoskeleton interacting protein, UNC-44, the sole homolog of mammalian ankyrin proteins, along with microtubule-interacting CRMP homolog, UNC-33, and an atypical kinesin, VAB-8, restricts the number of UNC-9 gap junction channels in PLM neurons; potentially by controlling the turnover of UNC-9 channels^22^. We have identified that while the UNC-44/UNC-33/VAB-8 pathway regulates the dynamics of UNC-7/UNC-9 heterotypic gap junction channels in PLM synapses, a distinct retrograde kinesin-14 family kinesin, KLP-15, specifically limits the number of INX-1 gap junction channels present within the same heterochannel synapses. KLP-15 possibly regulates the dynamics of INX-1 gap junction channels in association with other cytoskeleton-interacting proteins. We also identified that dynamic actin is required similarly for the localization of both INX-1 and UNC-7/UNC-9 gap junction channels within heterochannel electrical synapses in PLM neurons. It will require detailed analysis of the innexin protein domains, cytoskeleton-interacting proteins and putative innexin interacting proteins to understand the basis of innexin-specific mechanisms regulating electrical synapse dynamics. Our findings offer a unique opportunity to understand how distinct cell biological mechanisms interact with the F-actin network to independently regulate the dynamics of distinct gap junction channel types within an electrical synaptic plaque.

Building upon these, we further showed that the gap junction channel code of specific electrical synapses within a neuron is not static, but indeed can undergo reversible alteration when developing *C. elegans* larvae molts into the dauer diapause stage under harsh environmental conditions^17^. Specifically, INX-1 channels, but not UNC-7/UNC-9 channels, are selectively downregulated from the PLM-BDU synapses when the animal enters dauer stage, thereby altering the overall composition of the heterochannel synapse. Additionally, this change in the synaptic composition also correlates with the dauer stage specific reversible downregulation in the expression of INX-1-specific kinesin motor, *klp-4*. These findings illuminate the previous observation that the neuron-type specific expressions and splicing of individual innexin genes undergo widespread, combinatorial alteration during dauer stage, ultimately leading to substantial changes in the connectome and multiple aspects of dauer stage-specific behaviors^13^. Our results here show plastic changes in the composition of existing electrical synapses and concomitant changes in the expression of the molecular motor, which could be a generalizable mechanism to achieve synapse-level plasticity, ultimately affecting the function of the electrical synaptic circuit. Our findings provide a foundation for further nervous system-wide studies to understand the extent and functional implications of this novel mechanism of electrical synaptic plasticity across species.

## EXPERIMENTAL MODEL AND SUBJECT DETAILS

### *C. elegans* strains and handling

*C. elegans* strains were cultured at 20-22°C on NGM (nematode growth media) plates coated with *E. coli* (OP50) bacteria as a food source according to the standardized protocols unless mentioned otherwise^67^. *C. elegans* variety Bristol, strain N2 were used here as wild-type strain. We induced dauer arrest under starvation, crowding and high-temperature conditions, as described previously^13^. Dauer animals were selected by 1% SDS treatment.

Transgenic strains were generated using standard microinjection technique. 2 to 3 independent transgenic reporter lines were scored for each experiment. Injection criteria are listed in the strain list.

A complete list of strains and transgenes used in this study is listed in the Key Resources Table (Table S2).

## METHOD DETAIL

### Cloning and constructs

#### CRISPR/Cas9-mediated genome editing

Fluorescent knock-in alleles or deletion alleles were generated using purified Cas9 (IDT, catalog # 1081059), tracrRNA (IDT, catalog # 1072533) and crRNA (IDT) based on previously published protocols^68,69^. All gene specific crRNA sequences are listed in Table S1.

##### Generating deletion knockout for INX-1

A complete deletion allele of *inx-1, inx-1(amz11)* was generated using two guide RNAs flanking the gene locus. See schematics in Figure 1C.

##### Generating *inx-1::FLPon-gfp* reporter allele

FLPon-gfp tag was amplified from the *cla-1(wy1218)*^70^ and inserted into the endogenous *inx-1* locus, specifically tagging INX-1a at the C-terminus before the STOP-codon to generate *inx-1(amz08)*.

##### Generating inx-1::TagRFP-T and inx-18::eGFP reporter alleles

The loxP::tagRFP-T cassette was assembled by Gibson assembly (NEB). It was inserted in the C-terminus of the endogenous *inx-1* and *inx-18a* locus before the STOP-codon to get *inx-1(amz01)* and *inx-18(amz04)* alleles, respectively.

##### Generating *unc-7::gfp11X5* reporter allele

Referring to the GRASP method^71^, a gBlock DNA sequence having 5 tandem repeats of 11th beta-sheet of GFP containing GGSGG-liner sequence was obtained from Integrated DNA Technologies (IDT). This sequence was inserted at the C-terminus of endogenous *unc-7* locus before the STOP-codon to get *unc-7(amz15)* allele.

##### Generation of triple floxed *unc-7(amz14 ot895), inx-1(amz18), unc-9(amz32)* alleles for conditional knock-out experiments

We inserted a *loxP* site at the 5’ end of the *unc-7* genomic locus in the *unc-7*(*ot895[unc-7::tagRFP-T::loxP::3xFLAG*]) strain background (Figure 2K).

In the same background, we sequentially inserted two FRT-recombination sites flanking the *inx-1* gene locus, *inx-1(amz18)* (Figure 2K).

Continually, we inserted two FRT* recombination sites flanking the *unc-9* gene locus *unc-9(amz32)* (Figure 2K). Sequences for all the guide RNAs used are listed in Table S1.

###### Generation of TRN-specific *innexin* expression constructs

For TRN-specific expression of innexins *inx-1, inx-2, inx-10, inx-18, inx-7,* and *unc-9*, total cDNA was synthesized using oligo(dT)_20_ primer and SuperScript III First-Strand Synthesis System (Thermo Fisher Scientific; Catalog #18080051), according to the standard protocol. ORFs of specific innexin genes were amplified using gene-specific primers (Table S1), and cloned into pMiniT2.0 vector (NEB). Using Gibson assembly, these ORFs were subsequently cloned upstream of the TRN-specific cis-regulatory sequence *(mec-4p)* and downstream of either e*gfp* or *tagrfp-*T sequences followed by the *unc-54 3’UTR,* in the pPD95.75 backbone.

###### Generation of TRN-specific *unc-7b* expression construct

Genomic region encoding the *unc-7b/c* isoforms tagged to tagRFP-T at the C-terminus were amplified from the *unc-7(ot895[unc-7::tagrfp-t])* allele. Gibson assembly protocol (NEB) was used to assemble this amplicon together with TRN-specific *mec-4p* cis-regulatory element and the pPD95.75 vector backbone to get *mec-4p:: unc-7b::tagrfp-t::unc-7 3’UTR*.

###### Generation of Chimeric INX-1 protein with UNC-9 extracellular loops

Sequences corresponding to the *unc-9* extracellular loop (EC) 1 and 2, and *inx-1* transmembrane domain (TM) 2 and 3 were amplified from the *unc-9* and *inx-1* ORF clones, respectively. Using the PCR fusion approach^72^, a chimeric construct containing *unc-9* EC1 followed by *inx-1* TM2-TM3 followed by *unc-9* EC2, was generated. This chimeric sequence was subsequently inserted using Gibson assembly in the *mec-4p::inx-1ORF::egfp::unc-54 3’UTR* clone, in between *inx-1* TM1 and TM4, to generate: *mec-4p::INX-1TM1::UNC-9-EC1::INX-1-TM2-TM3::UNC-9-EC2::INX-1-TM4::egfp::unc-54 3’UTR* or *mec-4p::INX-1(*UNC-9 EC)::egfp::unc-54 3’UTR*.

###### Generation of PLM-specific *unc-9* rescue construct

*unc-9* ORF was amplified from total cDNA (as described earlier), using gene specific primers (Table S1). This was cloned downstream of TRN-specific *mec-4p* cis-regulatory sequence in the pPD95.75 vector (NEB), using Gibson assembly, to get *mec-4p::unc-9::unc-54 3’UTR* construct.

###### Generation of zebrafish Purkinje neuron-specific Cx43 and Cx43.4 expression vectors

zCCM22-Ca8-cfos-TRAP plasmid^25^ was used as backbone. Total cDNA from zebrafish larvae was synthesized using oligo(dT)_20_ primer and SuperScript III First-Strand Synthesis System (Thermo Fisher Scientific; Catalog #18080051), according to the standard protocol. Cx43 and Cx43.4 ORFs were amplified using gene-specific primers (Table S1) and cloned along with egfp ORF, using Gibson Assembly, to get zCCM22-Ca8-cFos::Cx43::egfp and zCCM22-Ca8-cFos::Cx43.4::egfp plasmids.

###### Identifying BDU-specific cis-regulatory element

Referring to single-cell transcriptomic data (CeNGEN)^12^, we identified three genes, *Y40H4A.2, F21C10.6* and *srh-165*, which show strong enrichment in BDU. Among them, a 1.4 kb upstream cis-regulatory region of only *Y40H4A.2* showed specific, but weak expression in BDU. Figure S1C shows schematics and coordinates of cis-regulatory regions used to generate reporter constructs with respect to the putative start codon of *Y40H4A.2*. Further refinement of the cis-regulatory region identified a smaller 430 bp region (prom6) capable of driving strong and reliable reporter gene expression in BDU (Figure S1C).

###### Generation of Cre, Flp and GFP(1-10) constructs

Sequences of TRN-specific driver, *mec-18p,* 3XNLS::Cre and T2A::ebfp2 were amplified using PCR and assembled with the pPD95.75 vector backbone, using Gibson assembly protocol (NEB) to get pAV07 (*mec-18p::3XNLS::Cre::T2A::ebfp2::unc-54-3’UTR*). Additional restriction enzyme sites between individual pieces were also introduced during PCR amplification steps for the ease of replacing individual components subsequently. To generate constructs for TRN-specific expression of Flp::2XNLS or GFP(1-10), 3XNLS::Cre sequence was replaced from the pAV07, using restriction enzyme sites (KpnI, NcoI and PstI). Sequences corresponding to BDU-specific cis-regulatory element, *Y40H4A.2p6* (Figure S1C) and PVC-specific cis-regulatory element, *nmr-1p* were amplified from the genomic DNA and cloned into *mec-18p::Flp* or *mec-18p::GFP1-10* construct, within SphI and SmaI restriction sites, to get BDU and PVC-specific Flp and GFP1-10 expression constructs.

### RNA interference by feeding

E. coli strain HT115 (DE3) producing dsRNA, specific to each target gene were used for the RNAi feeding experiment as per the standard protocol^73^. RNAi clones were cultured from single colonies overnight in LB broth containing 100 μg/ml carbenicillin and 10 μg/ml at 37°C. The NGM plates containing 100 μg/ml carbenicillin were coated with 6mM IPTG and seeded with the bacteria and grown at 37°C for 36hrs to induce the expression of dsRNAs. TRN (Touch Receptor Neuron)-specific gene knockdown was achieved by TRN-specific *sid-1* rescue in *sid-1(pk3321) him-5(e1490) V; lin-15B(n744) X; uIs71[myo-2p::mCherry; mec-18p::sid-1],* as described before^74^. RNAi sensitized L4s carrying the fluorophore-tagged innexins were placed onto the plates and grown at 22°C. F1s were empirically scored using Nikon Eclipse Ni-E microscope for the presence/absence or morphological differences of the innexin-puncta in PLM zone-1. Puncta phenotype associated with each gene knockdown was compared to the positive control. *inx-1*RNAi and *unc-9RNAi* was used as positive controls for respective screens. *gfp* RNAi or empty vector were used as the negative control. Candidate genes were selected based on their strong correlation with the positive control.

### Microscopy

Worms were anesthetized using 100mM of sodium azide and mounted on 5% agarose pads on glass slides. Worms were imaged using either Olympus FV3000 or Zeiss 980 confocal laser-scanning microscope with Airyscan. SIM images were captured using Zeiss Elyra 5 – Super-Resolution Microscope with lattice SIM. Image processing was done using LSM ZenBlue software, using SIM square processing mode. For Super-resolution imaging using SIM and Airyscan, worms were fixed in 2% PFA at RT for 5 minutes. RNAi-screening was done using Nikon NiE widefield fluorescent microscope with Hamamatsu ORCA fusion sCMOS camera. Images were analyzed by scanning the full Z stack using either Zeiss ZenBlue or Olympus Fluoview FV31S-SW software. Maximum intensity projections of representative images were constructed using NIH Fiji software. For DeActs experiments with inx-1, Nikon *Li* microscope fitted with an Olympus DP71 single-CCD colour camera with cellSens software was used. For DeActs experiments with unc-9, Olympus IX73 microscope with a 120W Photometrics Evolve 512 EMCCD camera was used. Figures were prepared using Adobe Photoshop CS7 and Adobe Illustrator CS7. Separate channels were usually adjusted independently using Levels and Curves in Adobe Photoshop.

### Automated worm tracking

Locomotion was tracked using an automated worm tracker system, WormLab (MBF Bioscience LLC, Williston, VT USA) at room temperature (22°C), To avoid potential variabilities, all strains that are part of a comparison were recorded simultaneously along with N2 wild-type. Minimal OP50 plates were made by coating 60mm NGM plates with 250 [l of 1:15 diluted overnight grown OP50 culture. The plates were allowed to dry in a laminar hood for 2 hours before starting the assays. Using an eyelash worm pick, young adult animals were placed on minimal OP50 plates, allowed to habituate for 5 minutes prior to recording. All videos were recorded at 7.5 fps. Duration of more than 20 frames (∼2.6 seconds) was used as threshold for categorizing it as a ‘reversal’ event. All analysis was performed using Wormlab software. The data was plotted using either GraphPad prism or by a python script (Figure 3C-D).

### Low threshold/gentle touch assays

Gentle touch assays using eye-lash were performed on young adult animals on NGM plates supplemented by minimal OP50 *E. coli* as described previously^8^. Scoring was done using five touches given to the tail before the anus, at about 3/4th length of the animal^75^. In order to maintain consistency, the animals with initial activity of moving forward or still were only assayed. Animals during their course of prolonged reversals, omega turns were not tested as they were shown to be non-responsive to mechanosensory stimulus^76^. Subsequent stimulus was provided only upon completing response to the previous touch or 1 second inter-stimulus gap was provided. For each trial, >30 animals were tested per genotype and the experiments were repeated in triplicates on different days.

### Co-expression of Connexins in zebrafish Purkinje neurons

Cx35b-expressing plasmid, zCCM22-Ca8-cFos-Gjd2b^53^, along with either zCCM22-Ca8-cFos-Cx43 or zCCM22-Ca8-cFos-Cx43.4 were co-injected in albino (nacre-/-) embryos at single-celled stage (at 30 ng/μL), along with 100 ng/μL Tol2 transposase mRNA^77^. Larvae were screened for sparse expression of both the tags in Purkinje neurons. At 4-7 dpf, the larvae were embedded in 1% low melting agarose for imaging.

### UNC-7 co-immunoprecipitation followed by LC-MS/MS

For co-immunoprecipitation (co-IP) experiments, *unc-7(ot895[unc-7::tagrfp-t::3XFLAG])* and *amzIs09[UPNp::tagrfp-T::3XFLAG]* were used as test and control strains, respectively. For protein extraction, 1.5 ml pellet of each genotype was resuspended in IP buffer [20 mM HEPES (pH7.5), 150 mM NaCl, 5 mM EDTA, 5 mM EGTA, 1 mM DTT, 3x Halt™ Protease Inhibitor Cocktail (Thermo Scientific, Cat. No. 78430), 1 mM sodium orthovanadate (Sigma-Aldrich, Cat. No. S6508), 20 mM β-glycerol phosphate (Sigma-Aldrich, Cat. No. 50020)] and were homogenized mechanically using disposable tissue grinders (Thermo Fisher Scientific). Homogenates were incubated overnight in IP buffer + 3% DDM (Anatrace, Cat. No. D310) at 4°C, followed by centrifugation at 15000 rpm at 4°C for 30 minutes. The resulting supernatant was used for co-IP. 9 mg of total protein extract was incubated with equilibrated ChromoTek RFP-Trap Magnetic Agarose beads (ChromoTek, Cat. No. rtma-20). IP-beads were washed thrice with 500µL of wash buffer containing increasing NaCl concentration (20 mM HEPES, pH 7.5, 150 mM – 500 mM NaCl, 5 mM EDTA, 5 mM EGTA, 0.02% DDM). Washed IP-beads were resuspended in 100 µL of 2xSDS-sample loading buffer and boiled for 5 minutes at 95°C. 60µL of the eluate was run on 8% SDS-PAGE gel and subjected to LC-MS/MS mass spectrometry for protein identification. LC-MS/MS was performed using Thermo Orbitrap Fusion and the raw files were analysed for peptide identification against the *C. elegans* UniProt database using Proteome Discoverer software (ThermoFisher Scientific). Data acquired from three biological replicates of both test and control strain was log-normalised to the total protein in each experiment and used for plotting (Figure 2J).

### Quantification and Statistical Analysis

Statistical tests were selected based on appropriate data type, distribution, and sample size. Across all statistical tests, p<0.05 was considered significant.

The Fisher-Freeman-Halton exact test, a generalization of Fisher’s exact test for contingency tables larger than 2×2, was used for categorical data, which includes all quantifications of innexin puncta counts in PLM synapses under different experiments, and the gentle touch-sensitivity assays, where group sizes were small and standard chi-squared approximations were not appropriate.

For continuous variables that include behavioural assays measuring backward time, forward time, idle time, mean amplitude, Shapiro–Wilk tests were first performed to assess normality. In cases where data were non-normally distributed and involved two-group comparisons, the Mann–Whitney U test (two-tailed) was used. As opposed to this, when data followed normal distributions, the Welch’s t-test was used for comparing means, not assuming equal SD in the individual groups. The use of both tests in the same plot was guided by genotype-specific distribution characteristics (described in figure legends). The Fisher’s exact test (2×2) was applied for binary outcome comparisons in quantifying percentage synaptic defects in (Figure 6).

## CONTACT FOR REAGENT AND RESOURCE SHARING

Further information, requests for resources and reagents should be directed to and will be fulfilled by the Lead Contact, Abhishek Bhattacharya.

## ACKNOWLEDGMENTS

We thank Arunima Sen, Rachna V, Marlyn Xavier Mascarenhas, Himani Dave, Supraja Balakrishnan, Sucheta Dey and Dipasmit Palchaudhuri for their help with the molecular biology and behavioural assays; Selvanayaki Eswaramoorthy for assistance in *C. elegans* injections; Souvik Modi in Koushika lab for subcloning DeActs for TRNs and making transgenic lines; wormbase.org and wormwiring.org for resources; Caenorhabditis Genetics Center (CGC), Kota Mizumoto and Anindya GhoshRoy for *C. elegans* strains; Raghu Padinjat, Kavita Babu, Oliver Hobert, Urmi Bandyopadyay, Hannes E. Buelow and members of the Bhattacharya lab for comments on this manuscript. This work was supported by India Alliance–Welcome-DBT (IA/I/20/2/505211) as well as intramural funds from National Centre for Biological Sciences – Tata Institute of Fundamental Research.

## AUTHOR CONTRIBUTIONS

A.V., S.M. and A.B. designed experiments and oversaw the project, A.V. designed and generated most of the strains, A.V. performed behavioral assays, S.M. performed kinesin RNAi-screens, mutant analysis and relevant strain generation, A. Bandyopadhyay did the UNC-7 Co-IP and mass-spec. S.V. did the zebrafish experiments, which V.T. supervised. N.V. did the DeActs experiments that S.P.K. supervised. A.V., S.M. and A.B. analyzed and interpreted data. A.V. and A.B. wrote the paper.

---

**TABLE S1.**
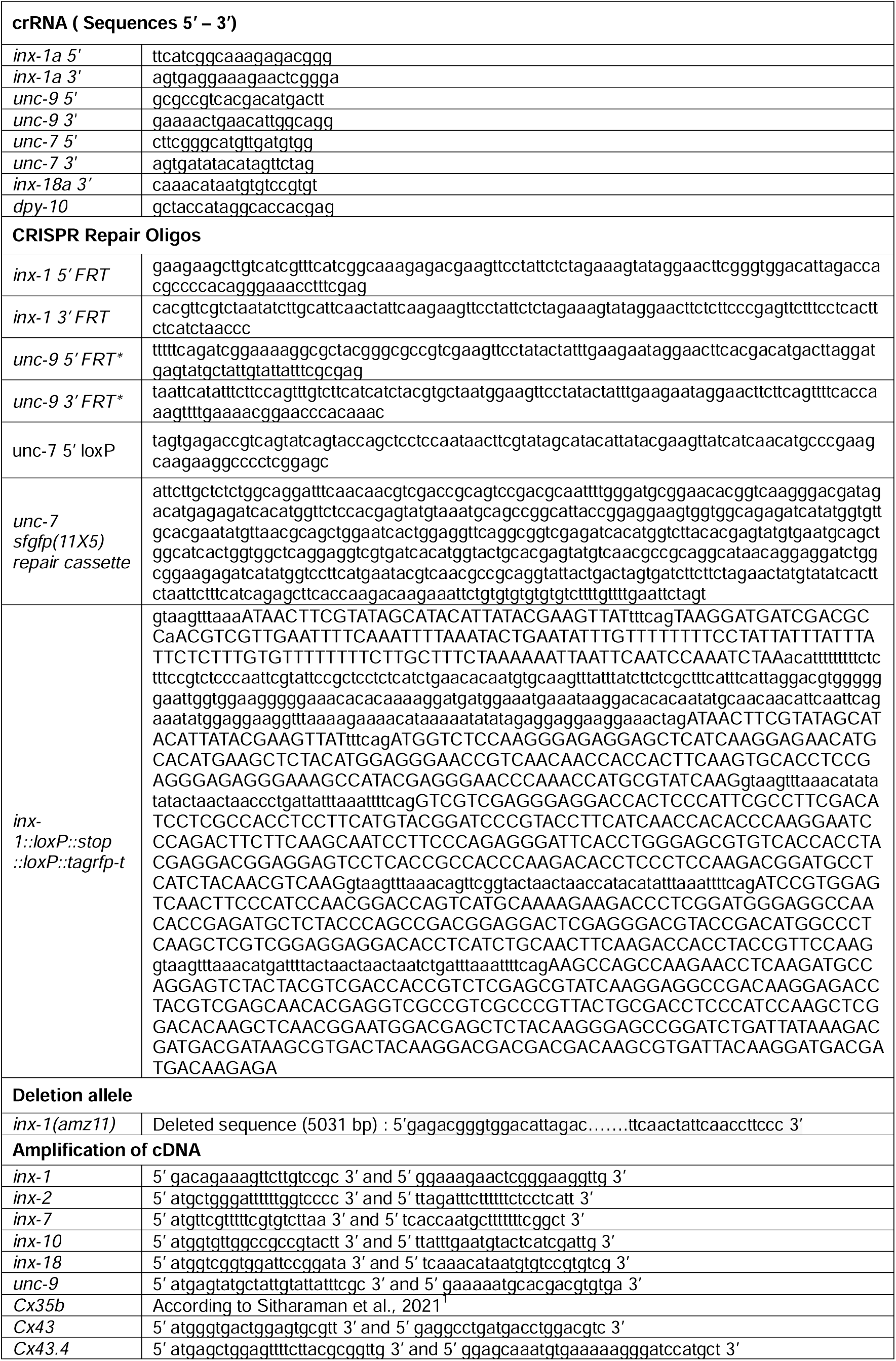
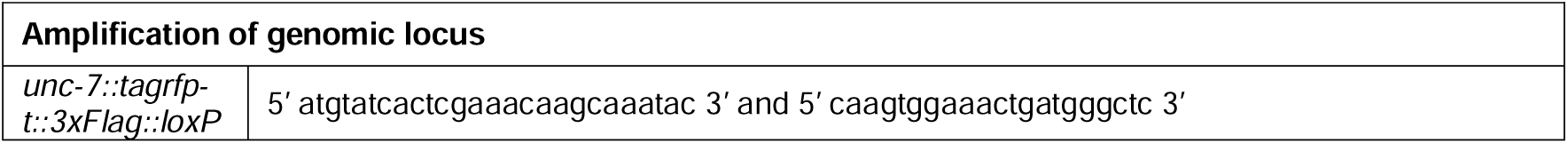
(Oligo list)

**TABLE S2.**
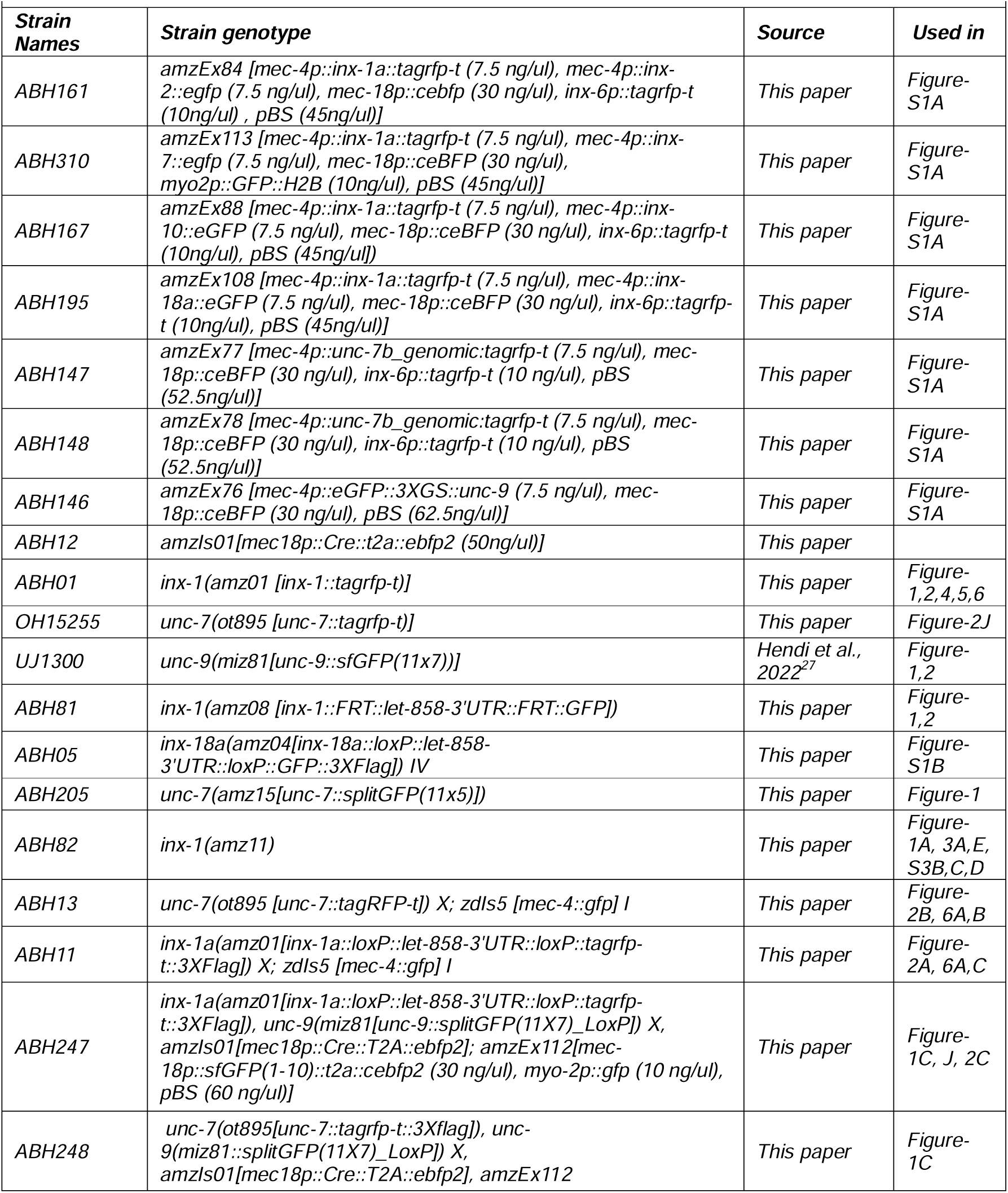
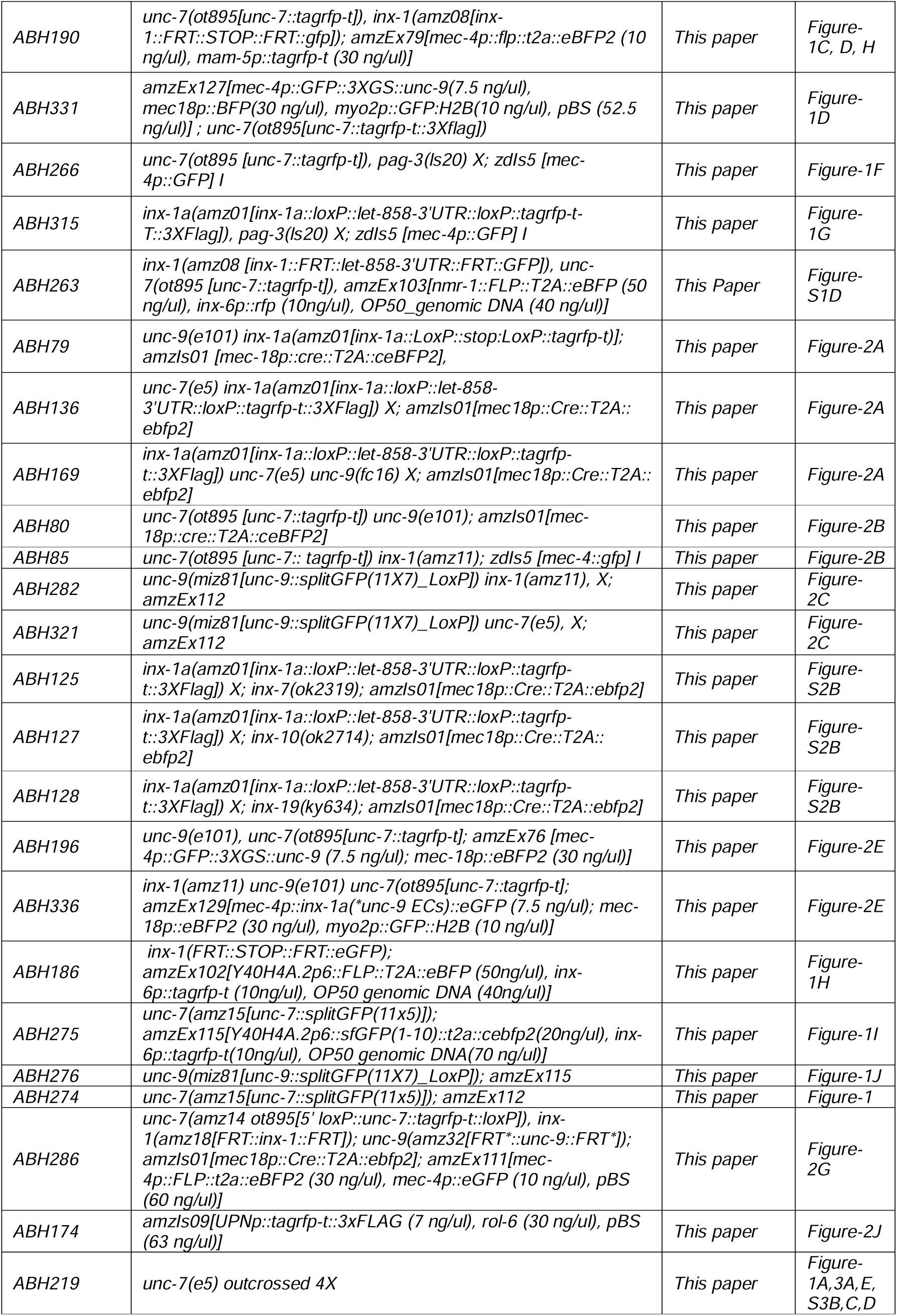
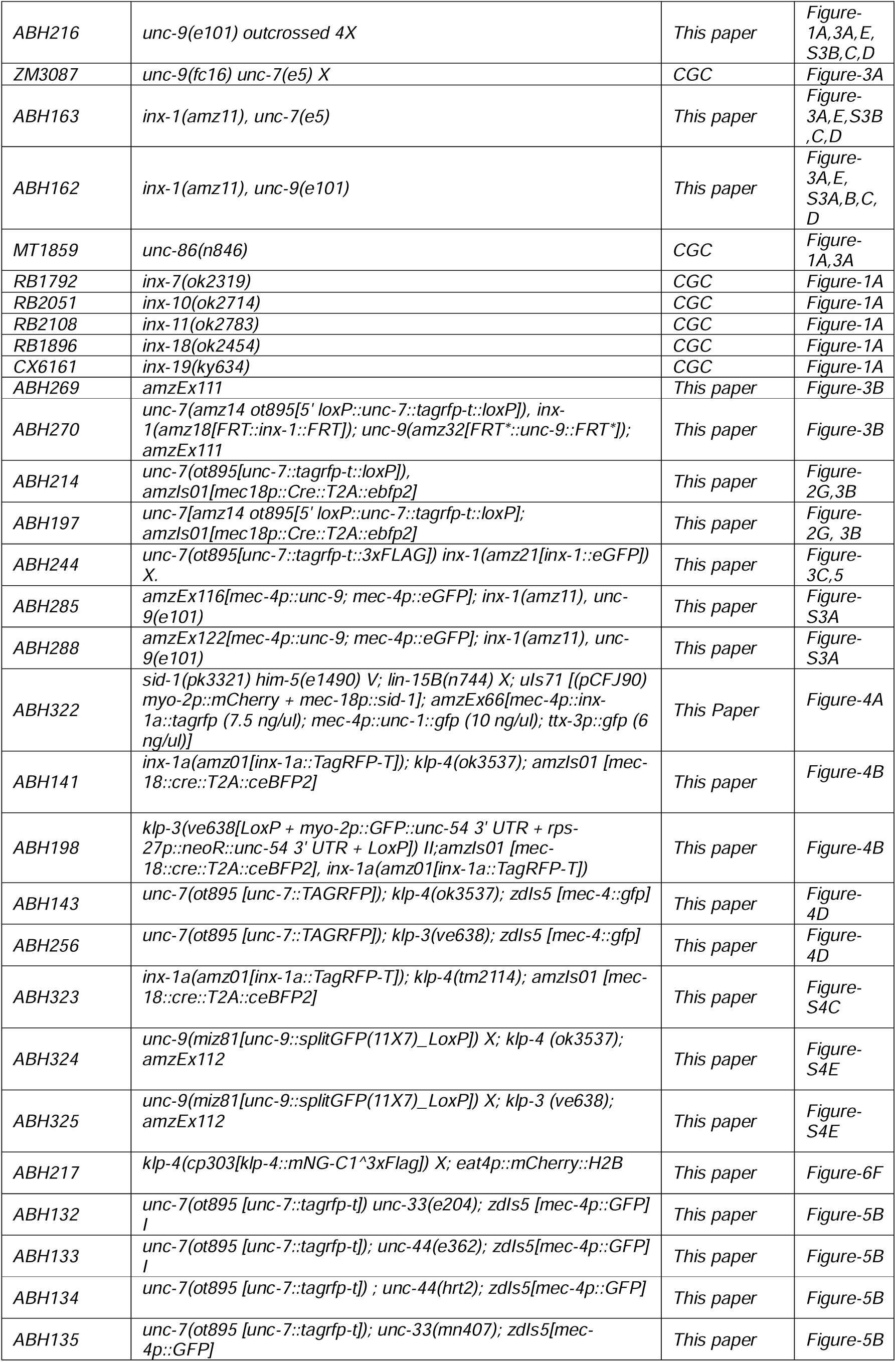
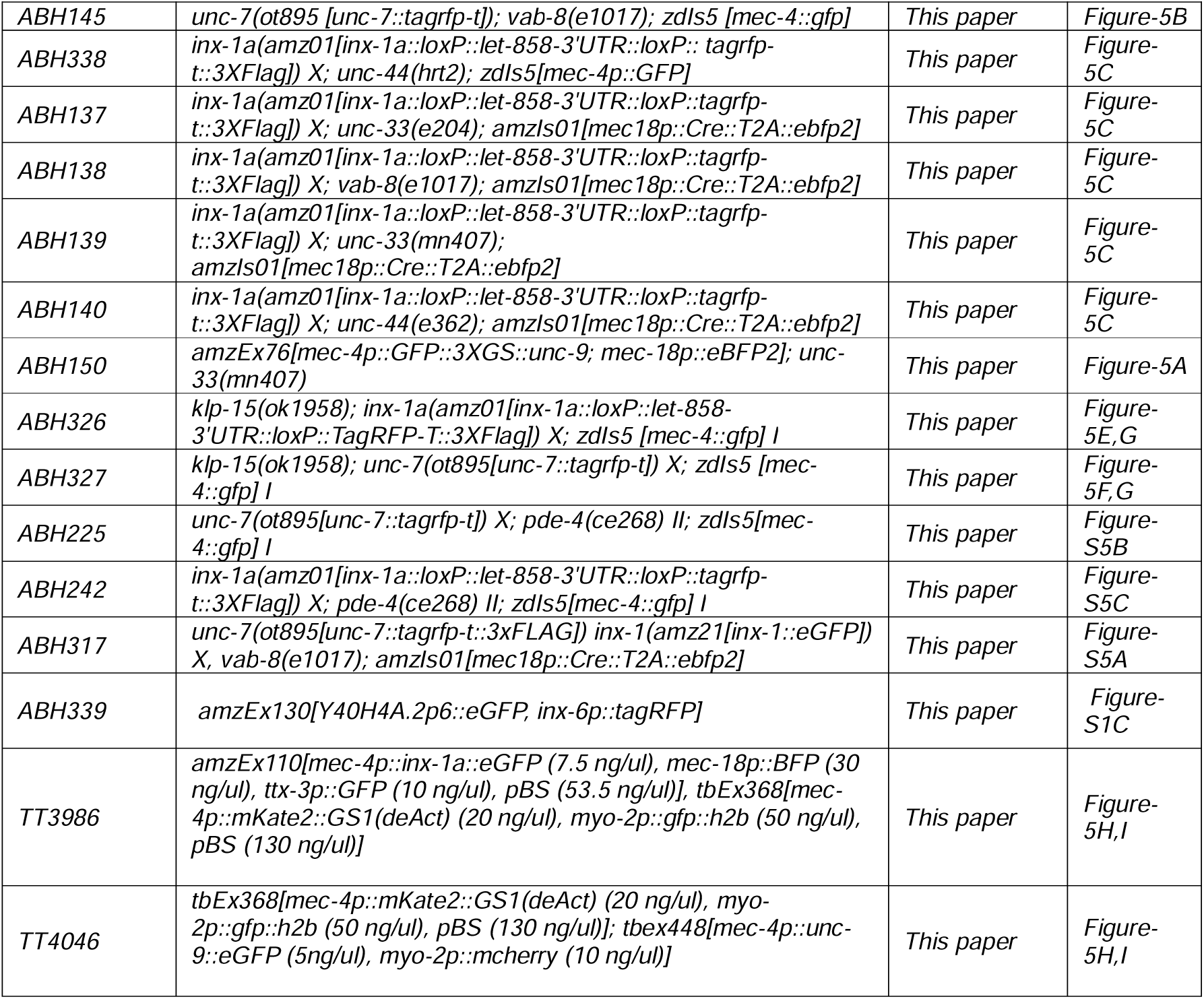
(Strain Index)

