## Supplementary figures and images for "Molecular Configuration, Regulation and Function of Heterochannel Electrical Synapses"

### Supplemental Figures

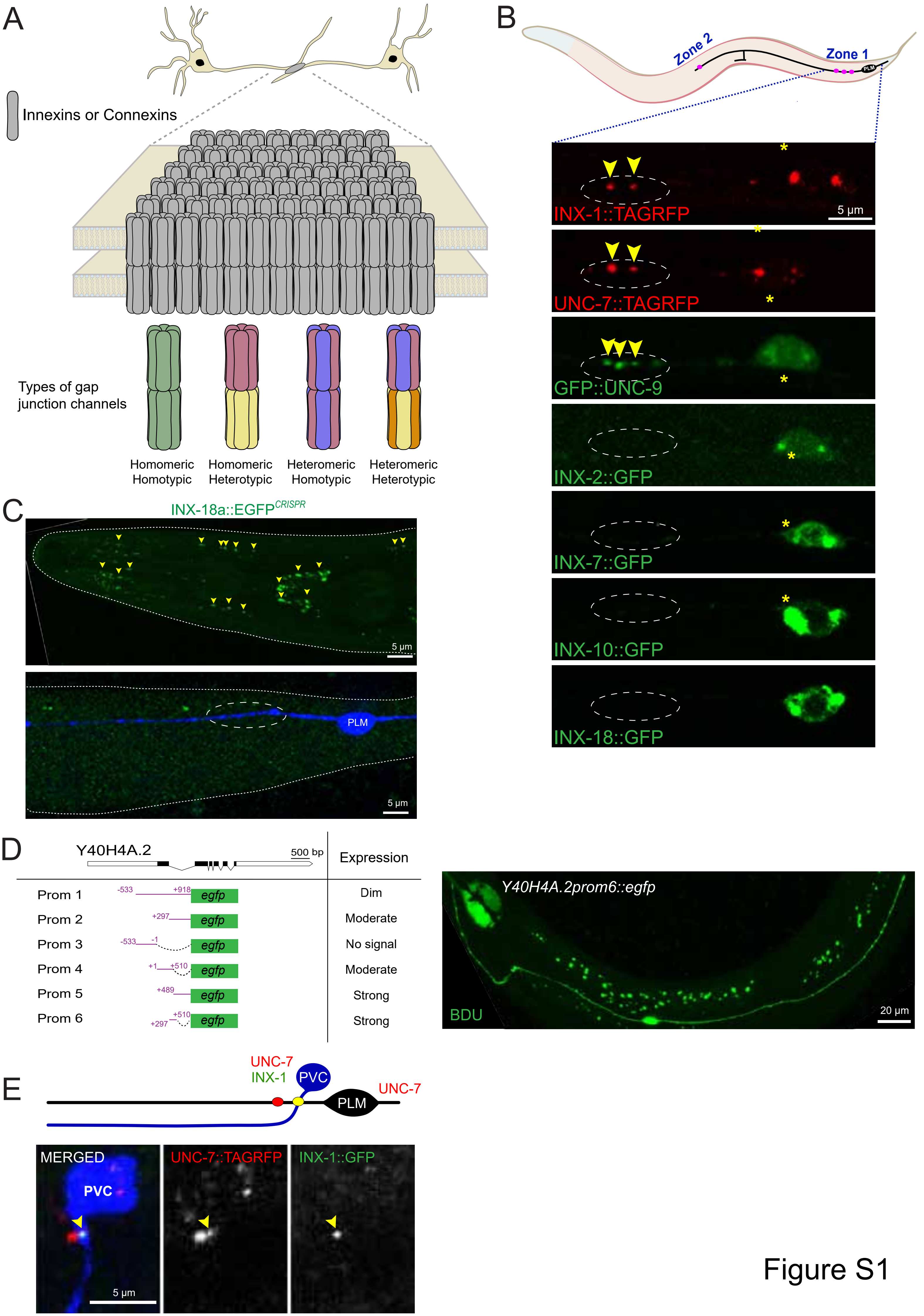

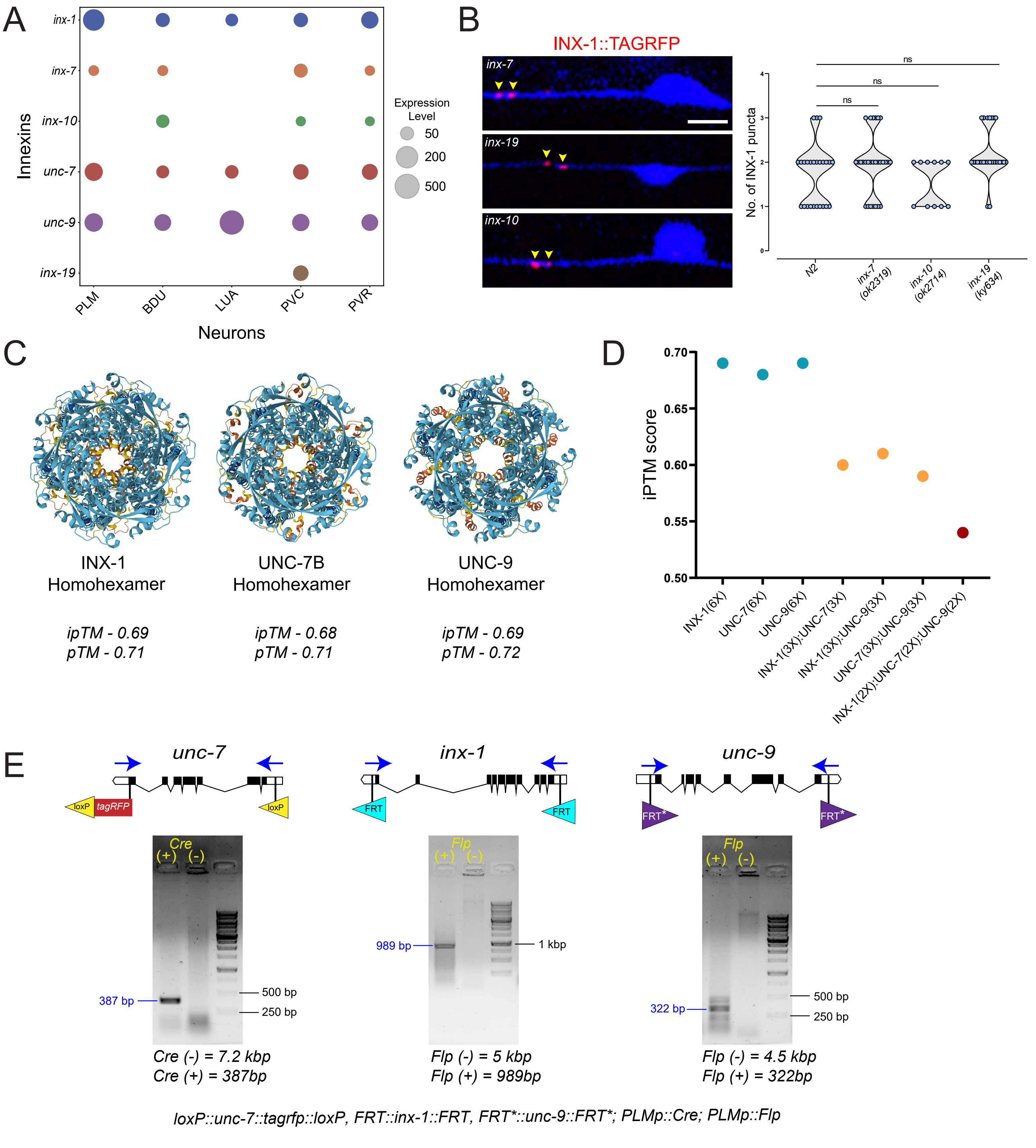

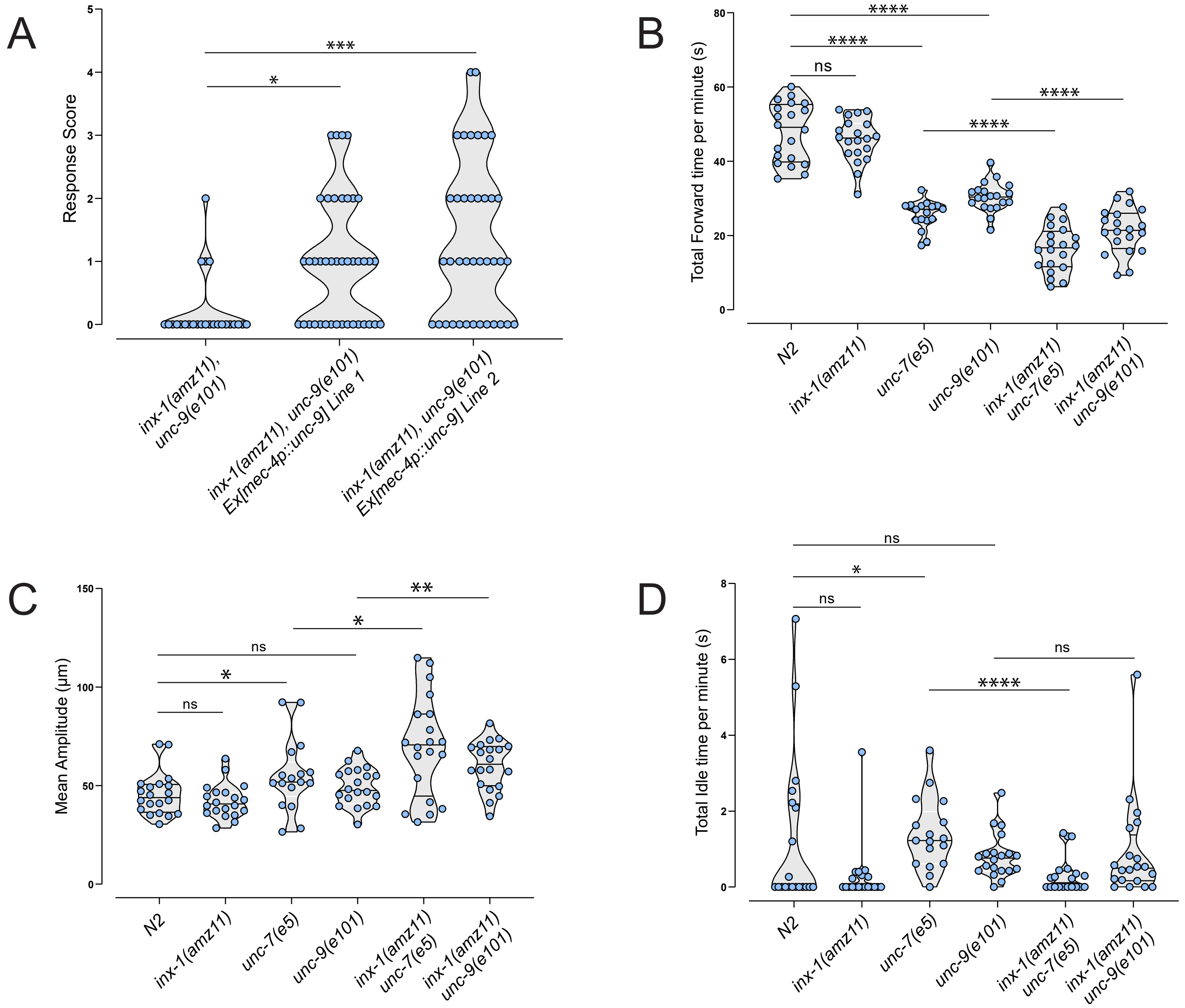

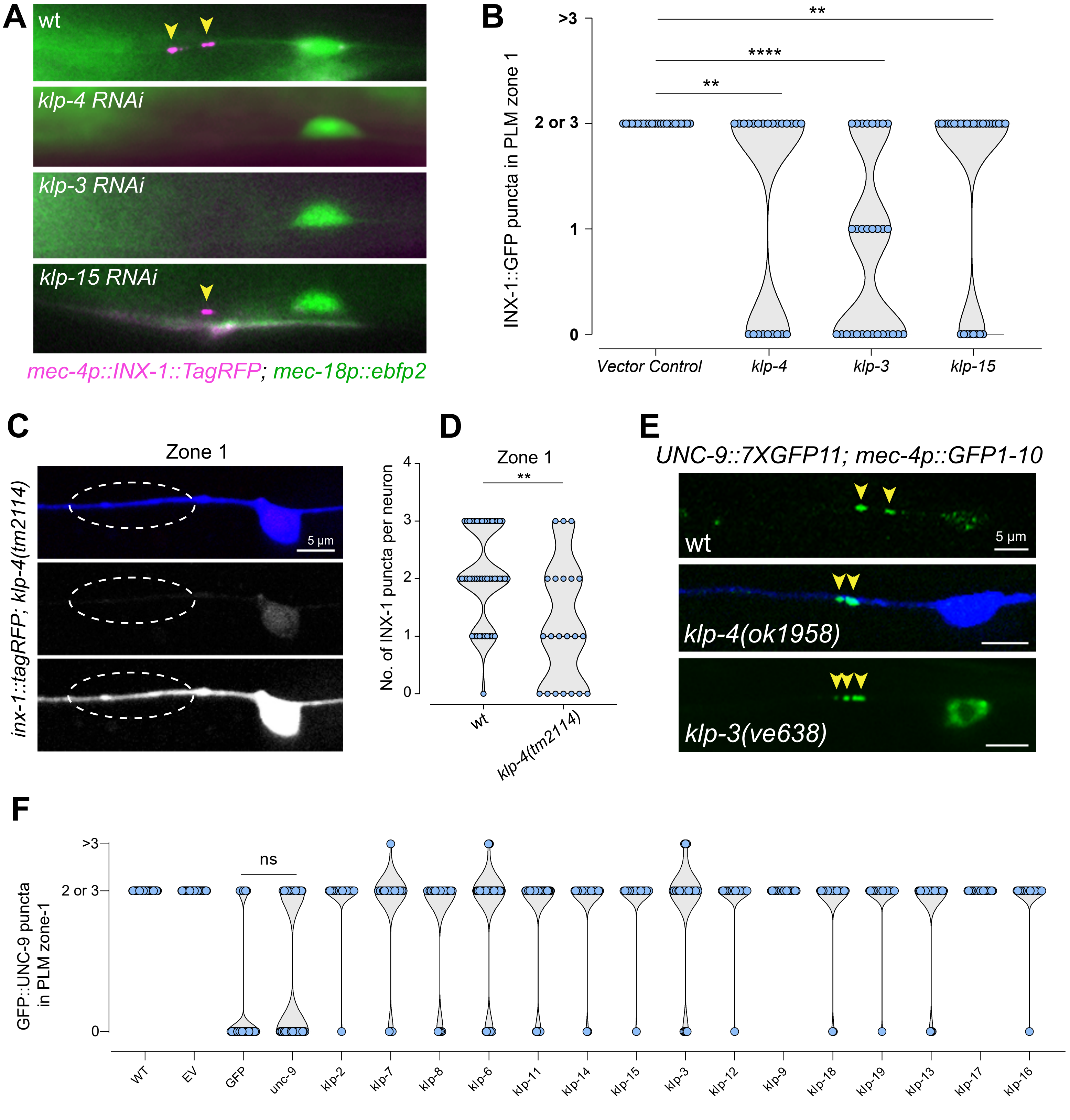

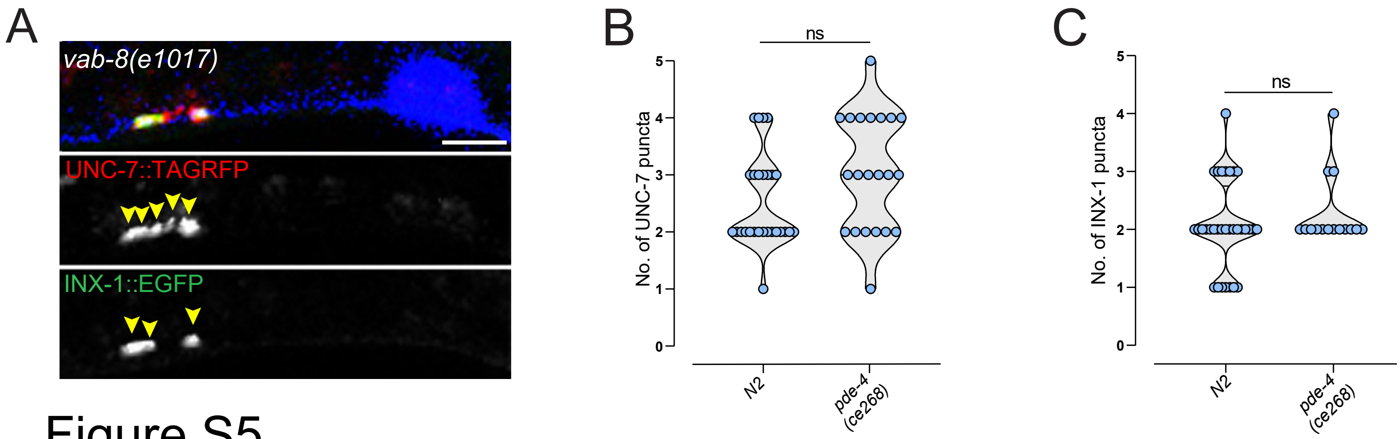
